# Maintaining a stable head direction representation in naturalistic visual environments

**DOI:** 10.1101/2022.05.17.492284

**Authors:** Hannah Haberkern, Shivam S Chitnis, Philip M Hubbard, Tobias Goulet, Ann M Hermundstad, Vivek Jayaraman

## Abstract

Many animals rely on a representation of head direction for flexible, goal-directed navigation. In insects, a compass-like head direction representation is maintained in a conserved brain region called the central complex. This head direction representation is updated by self-motion information and by tethering to sensory cues in the surroundings through a plasticity mechanism. However, under natural settings, some of these sensory cues may temporarily disappear—for example, when clouds hide the sun—and prominent landmarks at different distances from the insect may move across the animal’s field of view during translation, creating potential conflicts for a neural compass. We used two-photon calcium imaging in head-fixed Drosophila behaving in virtual reality to monitor the fly’s compass during navigation in immersive naturalistic environments with approachable local landmarks. We found that the fly’s compass remains stable even in these settings by tethering to available global cues, likely preserving the animal’s ability to perform compass-driven behaviors such as maintaining a constant heading.

## INTRODUCTION

Many animals are thought to rely on internal representations of their spatial relationship to their surroundings to perform flexible, goal-directed navigation (Tolman, 1948). Most spatial representations, including those of head direction (HD), are updated by self-motion information, but tend to drift without external sensory cues to tether to (Seelig and Jayaraman, 2015; Taube et al., 1990a, b). Even in the presence of external sensory cues, HD representations can be unstable in complex or ambiguous settings, for example, in surroundings with visual symmetries (Dan et al., 2021; Fisher et al., 2019; Jacob et al., 2017; Kim et al., 2019; Seelig and Jayaraman, 2015). Although such instabilities have been observed mainly in artificial visual settings, they raise questions of how HD systems perform in dynamic naturalistic environments. For example, in naturalistic settings, prominent cues, such as the sun, can be transiently obscured by clouds or vegetation (**Figure 1A**). Furthermore, terrestrial landmarks at different distances from an animal can change their angular position relative to the animal during translation, creating potential conflicts for a neural compass that is tethered to these cues (**Figure 1A,B**). In this study, we used visual virtual reality (VR) (Dombeck and Reiser, 2012; Haberkern et al., 2019; Kaushik et al., 2020) (**Figure 1B,C**) to explore whether and how the HD representation of the fly, *Drosophila melanogaster*, copes with such challenges, and how this impacts the fly’s behavior.

**Figure 1:**
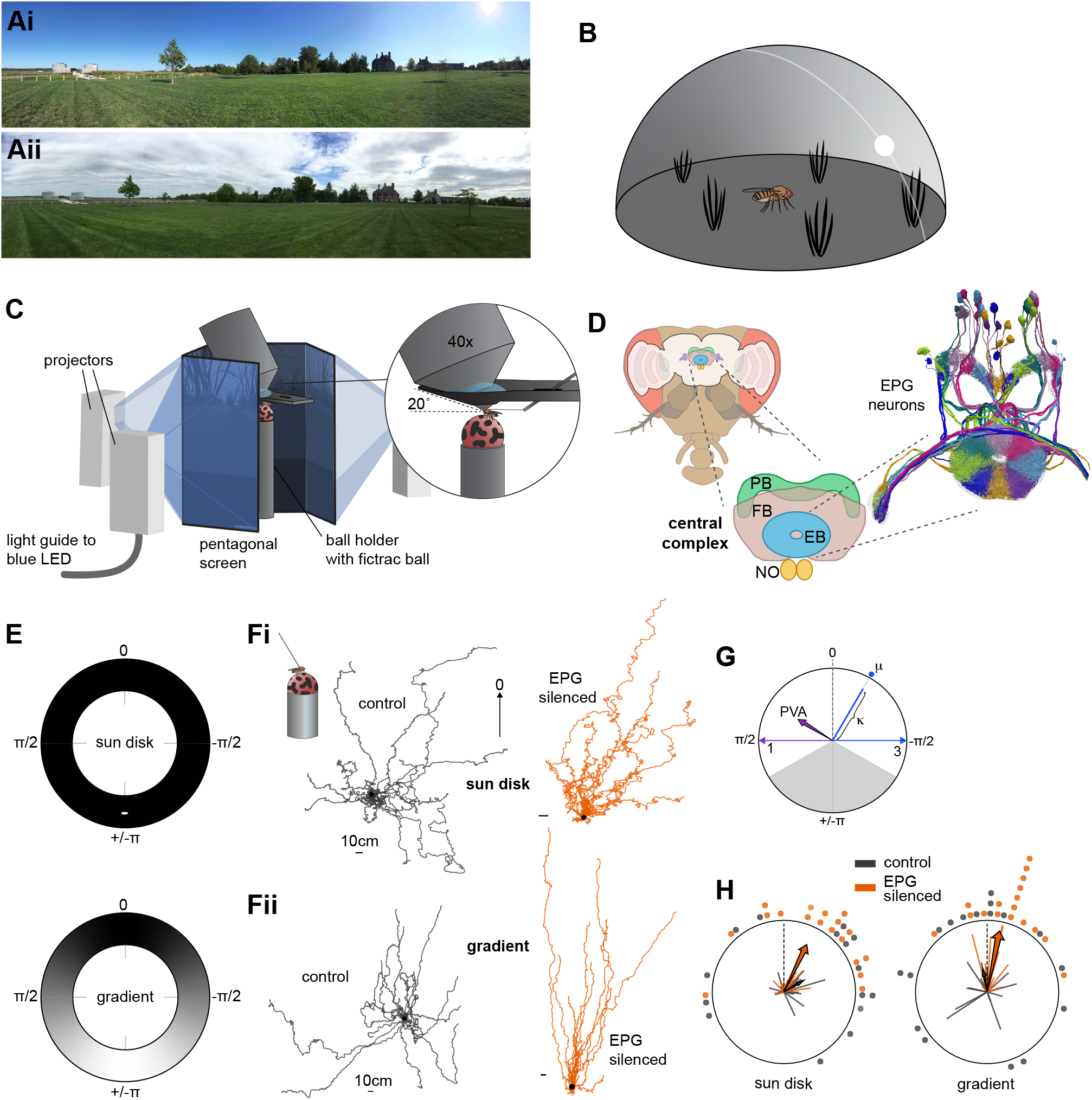
Flies perform menotaxis in sky-like visual virtual reality environments, but only with an intact neural compass. **A:** Panoramic photographs showing the same scene with a clear sky (Ai) or with partial cloud cover (Aii). **B:** Schematic of simulated visual stimuli: celestial cues such as a sun disk and intensity gradient, and terrestrial grass tufts. **C:** Illustration of the immersive, projector-based virtual reality system used in imaging experiments. The inset shows occlusions of the field of view by the holder. **D:** Schematic of the fly brain within the head, and within the brain the central complex with the four main neuropils. PB: protocerebral bridge, FB: fan-shaped body, EB: ellipsoid body, NO: noduli. Right: Rendering of an electron-microscopy reconstruction of EPG neurons. Different neurons are colored in a way to highlight the tiling of arbors around the EB. **E:** Illustration of chosen reference frames for sun disk (left) and gradient (right) panorama environments. **F:** Virtual trajectories of control flies (SS00096 > WTB, left, n=20, black trajectories) and flies with EPG neurons silenced (SS00096 > Kir, right, n=20, orange trajectories) in the sun disk environment (Fi) or intensity gradient environment (Fii). The black dot marks the starting point. Scale bars: 10 cm. **G:** Explanation of fixation plots in H. Spines (radial lines) indicate the fitted μ and k values of head direction distributions from single trials (one per fly). The μ values use the reference frame convention indicated in A. The radial axis for k values spans 0 to 3. The arrows indicate the population vector average (PVA) of all spines in a group. The radial axis of the PVA vector spans 0 to 1. **H:** Fixation plots for control flies (SS00096 > WTB, left, n=20) and flies with EPG neurons silenced (SS00096 > Kir, right, n=20) in the sun disk (left) and gradient (right) environment.

A highly conserved insect brain region called the central complex (CX) (**Figure 1D**) has been implicated in a wide range of goal-directed navigational behaviors (Dan et al., 2021; el Jundi et al., 2015; Giraldo et al., 2018; Green et al., 2019; Kuntz et al., 2017; Neuser et al., 2008; Ofstad et al., 2011; Strauss and Heisenberg, 1993; Triphan et al., 2012; Turner-Evans et al., 2021). The CX generates and maintains the insect HD representation (Green and Maimon, 2018; Homberg, 2004; Honkanen et al., 2019; Hulse and Jayaraman, 2019; Pfeiffer and Homberg, 2014; Turner-Evans and Jayaraman, 2016). In flies, the HD representation is maintained in a recurrent network that includes a population of EPG or compass neurons that each innervate individual wedges of the toroidal ellipsoid body (EB), a substructure of the CX (Fisher et al., 2019; Green et al., 2017; Kim et al., 2019; Kim et al., 2017; Seelig and Jayaraman, 2015; Turner-Evans et al., 2017; Turner-Evans et al., 2021; Wolff et al., 2015) (**Figure 1D**).

The EPG population receives information about localizing sensory cues through synaptic inputs from so-called ring neurons (Fisher et al., 2019; Hardcastle et al., 2021; Hulse et al., 2021; Kim et al., 2019; Okubo et al., 2020; Omoto et al., 2017; Seelig and Jayaraman, 2013; Turner-Evans et al., 2021). Plasticity at this connection is thought to enable the relative positions of diverse sensory cues to be mapped onto the compass (Cope et al., 2017; Dan et al., 2021; Fisher et al., 2019; Green and Maimon, 2018; Kim et al., 2019). Ring neurons bring multimodal inputs to the EPG neurons (Hardcastle et al., 2021; Okubo et al., 2020; Omoto et al., 2017; Seelig and Jayaraman, 2013; Shiozaki and Kazama, 2017; Sun et al., 2017), but our focus here was on visual information. Across insects, the anterior visual pathway (AVP) brings information about polarized light e-vector orientation and visual features from the optic lobe to the EB via the anterior optic tubercle (AOTu) and bulb (BU) (el Jundi et al., 2014a; Hardcastle et al., 2021; Heinze et al., 2009; Held et al., 2016; Mappes and Homberg, 2007; Omoto et al., 2017; Pegel et al., 2018; Sun et al., 2017; Trager et al., 2008; Vitzthum et al., 2002; Zittrell et al., 2020). In flies, most visually tuned ring neurons display spatiotemporal responses to visual features, with most showing strongly ipsilateral receptive fields (Hardcastle et al., 2021; Omoto et al., 2017; Seelig and Jayaraman, 2013; Shiozaki and Kazama, 2017; Sun et al., 2017). Thus, a given visual environment evokes a characteristic pattern of activity in visually responsive ring neurons that provide their input to the EPG population. Consistent with such visual inputs, previous work has shown that the fly’s compass generates a stable HD representation—as judged by the stability of the dynamics of its EPG population—in a variety of different visual settings ranging from those with a single sun-like disk or vertical bar to more complex scenes with multiple visual cues (Kim et al., 2019; Seelig and Jayaraman, 2015). EPG neurons and their visual inputs are required for flying and walking flies to maintain a specific bearing relative to a prominent cue in their surroundings, a behavior called menotaxis (Giraldo et al., 2018; Green et al., 2019; Turner-Evans et al., 2021). However, how the compass system and the fly itself respond in more naturalistic visual settings is less explored.

Our focus in this study was on the flies’ ability to maintain stable head direction estimates across changing celestial cues and when navigating through terrestrial objects like grass. We focused on two types of visual celestial cues that are known to be used by insects to orient: a sun-like disk and an intensity gradient (el Jundi et al., 2014b; Ugolini et al., 2009; Warren et al., 2018). A localized sun disk cue is known to support a stable head direction estimate, as measured by EPG calcium responses (Giraldo et al., 2018), but we asked if a widefield intensity gradient could also serve as a compass cue. In a natural sky these two celestial cues occur in a specific relationship to each other, with the brightest part of the intensity gradient centered at the sun. So, we asked whether the compass and the fly would respond consistently regardless of which of these cues they were provided with, and whether being primed by a sun disk might recruit the plasticity of the compass circuit (Fisher et al., 2019; Kim et al., 2019) to more firmly tether the compass to the intensity gradient. Finally, we assessed how well flies were able to orient when terrestrial grass tussocks transiently obscured parts of the virtual sky. We found that flies used the sun disk and intensity gradients almost interchangeably, and that, as long as they had these global cues and an intact compass network, they maintained stable HD dynamics and were able to maintain their bearings even in settings with local clutter.

## RESULTS

We sought to probe the fly’s ability to maintain its bearings in visual settings that capture characteristics of natural scenes with different celestial compass cues—a sun-like disk or an intensity gradient (Fig 1A,E)—as well as approachable terrestrial objects that can occlude the celestial cues (Fig 1B). To test the stability of the Drosophila compass under such conditions (**Figure 1B**), we built a modular virtual reality setup based on the Unity game engine, which allowed head-fixed flies to explore an immersive visual environment while walking on an airsupported trackball (Haberkern et al., 2019; Moore et al., 2014; Seelig et al., 2010) (**Figure 1C**; see also **Methods** and **Figure 1—Figure Supplements 1-4**). We installed our VR system under a two-photon laser scanning microscope to enable calcium imaging while the fly explored a virtual open field with different skylight cues and also set up a replica of our VR system in a purely behavioral setup (**Figure 1B, Figure 1—Figure Supplement 1**).

We first asked how flies would respond behaviorally to a challenge faced by any diurnal animal that uses the position of the sun for navigation: the sun disappearing behind clouds (illustrated in **Figure 1A**). Specifically, we asked whether walking flies would perform menotaxis with a sunlike disk to help them get their bearings in our VR setup, and whether they would also display this behavior with a widefield intensity gradient simulating the azimuthal intensity changes across the sky (**Figure 1A,E**). Behavioral studies focused on walking flies have typically used vertical stripes rather than sun-like disks as cues, finding that flies frontally fixate on high-contrast stripes and otherwise perform menotaxis, maintaining a specific heading relative to the stripe (Green et al., 2019; Haberkern et al., 2019; Horn and Wehner, 1975; Strauss and Pichler, 1998; Turner-Evans et al., 2021). We found that flies performed menotaxis in both visual conditions we presented them with, displaying individualized heading preferences relative to the sun disk (**Figure 1Fi** left, **H**) and to the peak of the intensity gradient (**Figure 1Fii** left, **H**).

Menotaxis during both flight and walking is known to involve EPG neurons (Giraldo et al., 2018; Green et al., 2019; Turner-Evans et al., 2021) (**Figure 1D**). We expressed an inwardly rectifying potassium channel, Kir2.1, in these neurons (see **Methods**) to silence them and assess how flies behaved in our visual conditions without their EPG-based compass. With the EPG neurons silenced, we found that flies were no longer able to display individual heading preferences (**Figure 1F** right, **H**). Instead all flies chose to walk directly away from the disk, displaying something akin to a behavior previously reported to occur during flight (Maimon et al., 2008) (**Figure 1Fi** right). Similarly, in the intensity gradient, all flies displayed negative phototaxis, walking directly away from the brightest part of the gradient (**Figure 1Fii**, right). Note that the trajectories are straighter for flies walking in the intensity gradient than with the sun disk, potentially because the small sun can disappear behind the fly, leaving the fly temporarily in the dark with no visual information about its bearing. We conclude that walking flies can use either a sun-like cue or a widefield intensity gradient to maintain individualized headings within their surroundings, and that this behavior depends on an intact EPG-based compass system.

Having confirmed that menotaxis in our setup requires the EPG neurons, we next targeted them for imaging with the genetically encoded calcium indicator, GCaMP7f (Dana et al., 2019), focusing in particular on EPG processes in the EB (Hanesch et al., 1989b; Hulse et al., 2021; Lin et al., 2013; Seelig and Jayaraman, 2015; Turner-Evans et al., 2021; Wolff et al., 2015) (**Figure 2A**). As reported before, EPG population activity is organized into a bump that moves around the EB tracking the fly’s turns on the ball and, in visual settings, the movement of visual cues around the fly (Fisher et al., 2019; Kim et al., 2019; Seelig and Jayaraman, 2015) (**Figure 2B,D**). In previous work, compass dynamics have been characterized by computing an EPG population vector average that is tracked over time (Kim et al., 2019; Seelig and Jayaraman, 2015). This approach relies on there being a single activity bump in the EPG population. However, in some visual settings—for instance, those with visual symmetries or similar visual features at different angular orientations—EPG population activity can fluctuate between different offsets, sometimes transiently producing multiple competing bumps (Dan et al., 2021) (**Figure 2C-E**). In these conditions, the angular offset between the bump and the surrounding world can also change (**Figure 2C-E**, seconds 75-90). To account for multiple, temporarily coexisting bumps of activity, and different transiently stable offsets, we computed the bump position as the peak of the calcium activity and tracked the relationship of each peak relative to the animal’s orientation in the world (Dan et al., 2021) (**Figure 2C-F, Figure 2—Figure Supplement 1**). Over the course of a trial, EPG population activity sometimes switched offsets, but we found that one offset typically dominated. Thus, in this manuscript, we assessed the stability of compass dynamics by: (a) examining the number of different offsets present, (b) the fraction of time that the primary offset dominates, and (c) the reliability of the primary offset over the course of a trial (**Figure 2D-G;** see **Methods**).

**Figure 2:**
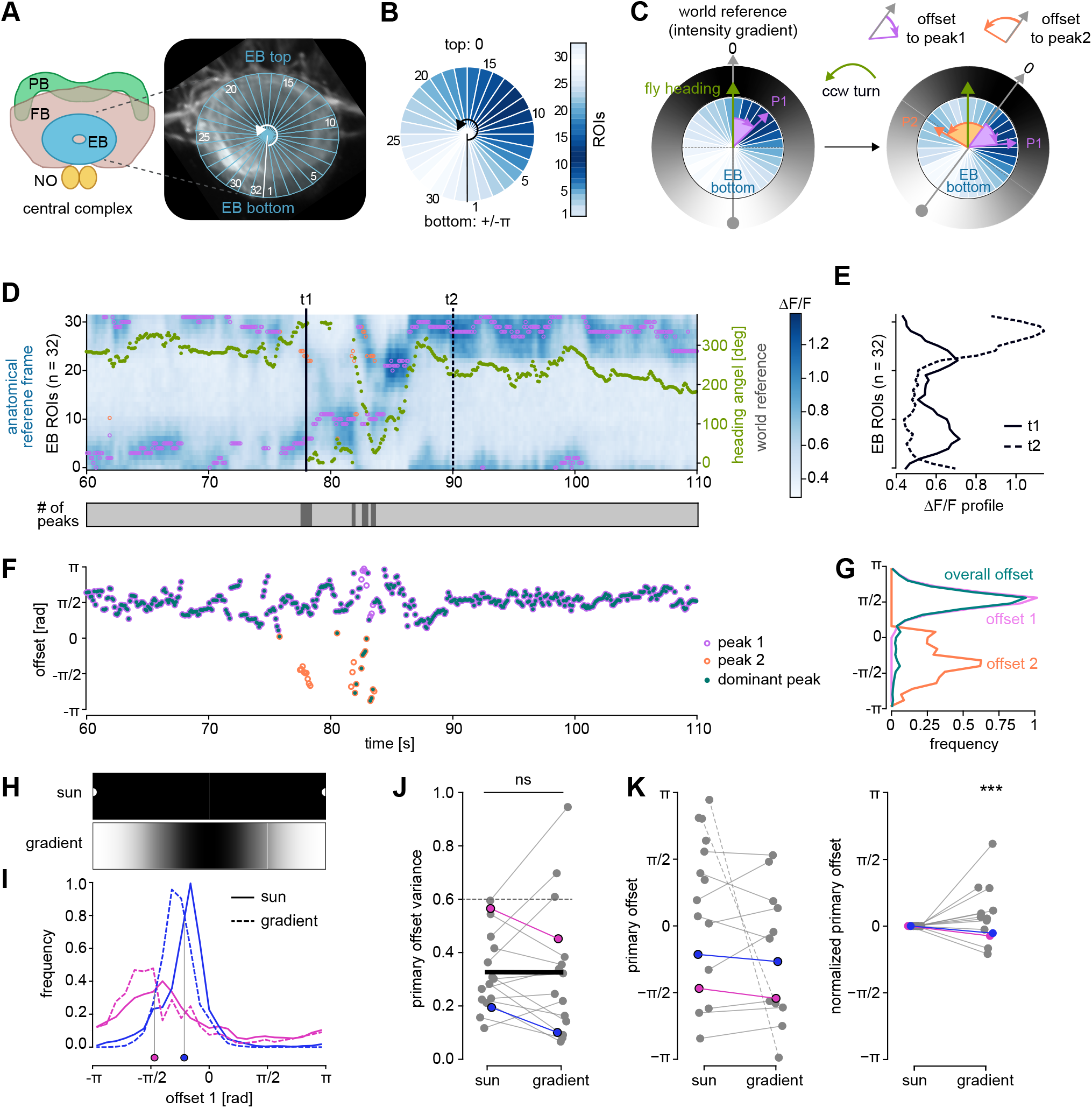
Fly compass dynamics are stable when flies walk under virtual skies with either sun-like cues or intensity gradients. **A:** Illustration of the CX on the right, and an average ΔF/F image from a calcium imaging experiment of EPG neurons expressing GCaMP7f on the left. Overlaid on the ΔF/F image is the segmentation of the ellipsoid body (EB) into 32 regions of interest (ROIs). **B:** Visualization of ΔF/F across the ROIs using a color-code. For each time frame, a vector is generated that captures the calcium activity across the 32 ROIs, which is used to visualize the position of the calcium activity bump in compass neurons within the EB over time (see D). **C:** Illustration of the computation of offsets between the fly’s neural compass and the external environment using two snapshots of EPG activity in the EB. The offset is computed as the angle (curved purple arrow) between the position of peak EPG activity in the EB (P1, straight purple arrow) and the darkest part of the visual surroundings (gray arrow). The fly’s heading relative to the external environment is indicated by the green arrow. When the fly turns (note that the green arrow points at a different part of the visual scene in the two snapshots), the offset is usually preserved. Occasionally, there is a transient second peak in EPG activity (P2, straight orange arrow), which is used to compute a second offset (curved orange arrow). Snapshots of EPG activity shown are from a SS00096 > GCaMP7 fly walking inside an intensity gradient world. **D: (top)** Sample trial showing ΔF/F of the 32 EB ROIs over time, with the ΔF/F values color-coded in blue. Pink and orange circles mark peaks in the ΔF/F distribution for each frame. Green dots mark the head direction within the virtual world (axis on the right). **(bottom)** The number of peaks in EPG activity that were detected in each frame (white: 0, light grey: 1, dark grey: 2). **E:** Distribution of ΔF/F across ROIs at two time points, t1 and t2, indicated in D by two black vertical lines. Note two peaks at t1. **F:** Time series of the offset angles computed for each ΔF/F peak (pink and orange circles). The offset value quantifies the relationship between calcium activity bump position (measured as ΔF/F peak location) in the EB and the head direction angle. As multiple peaks are possible, multiple offset values are possible. The overall offset is chosen as the one corresponding to the highest ΔF/F peak at a given time point. **G:** Distribution of offset angles for the two offset candidates (pink and orange), and the overall offset (teal line). The pink offset candidate here corresponds to the primary offset. **H:** Sun disk and gradient panoramas aligned to the shared world reference frame where +/-π aligns with the bright part of the panorama (see **Figure 1 Supplement 2** for reference). **I:** Frequency distributions of offset values for two flies (SS00096 > GCaMP7). Solid lines show data from sun disk trials, dashed lines from gradient trials. The grey lines indicate the circular mean of the sun trial offset distributions (offset location values). **J:** Circular variance of the primary offset distribution of SS00096 > GCaMP7 flies (n=16) from sun and gradient trials. Data points from the same fly are connected by a line. Data points from the two flies shown in (I) are colored in blue and pink respectively. The black line connects the two population means. The variance does not differ significantly between the two conditions (paired t-test, statistic=0.03071, p=0.9759). **K:** Same as J but for the primary offset (left) and the normalized primary offset (right). The connecting lines for two flies are shown as dashed, as they appear further apart on this visualization than the actual circular distance between them. The primary offset was normalized by subtracting the primary offset in the sun disk trial from the offset in both trials for each fly. Only data points where the offset variance was below 0.6 are shown (see J). For the normalized primary offset in the gradient trial (right) the points were non-uniformly distributed (Rayleigh test, statistic = 0.1967, p = 0.00002 (***)), indicating that the offset does not change significantly between these conditions.

We next sought to assess the stability of compass dynamics in these two environments (**Figure 2H**). Considering that ring neuron receptive fields are large (Seelig and Jayaraman, 2013; Sun et al., 2017), we hypothesized that widefield intensity cues alone could suffice for the fly compass to function stably. This was indeed the case: the variability around the primary offset did not differ significantly between the sun and gradient environment (**Figure 2I, J**, see also **Figure 2—Figure Supplement 1**). Importantly, the offset itself was largely preserved across the two environments (**Figure 2K**), implying that the fly’s internal representation of head direction was preserved when transitioning from the sun disk to the intensity gradient setting.

Previous studies have revealed plasticity mechanisms that enable visual cues to be rapidly mapped onto the fly’s compass (Fisher et al., 2019; Kim et al., 2019), meaning that exposure to a particular scene can influence how the compass tethers to another. We reasoned that if some ring neurons respond to both sun disks and intensity gradients, as suggested by the results from **Figure 2**, the plasticity mechanisms might enable prior exposure to a sun disk to ‘prime’ the compass to tether more stably to different intensity gradients as well. We tested this in two steps. First, to assess the robustness of the compass system to different strengths of intensity gradients, we exposed flies to three additional gradient conditions of lower contrast and one of higher average brightness (**Figure 3A**, **Figure 1—Figure Supplement 2B**). We found that the compass became less stable in environments where the intensity gradient was weaker.

**Figure 3:**
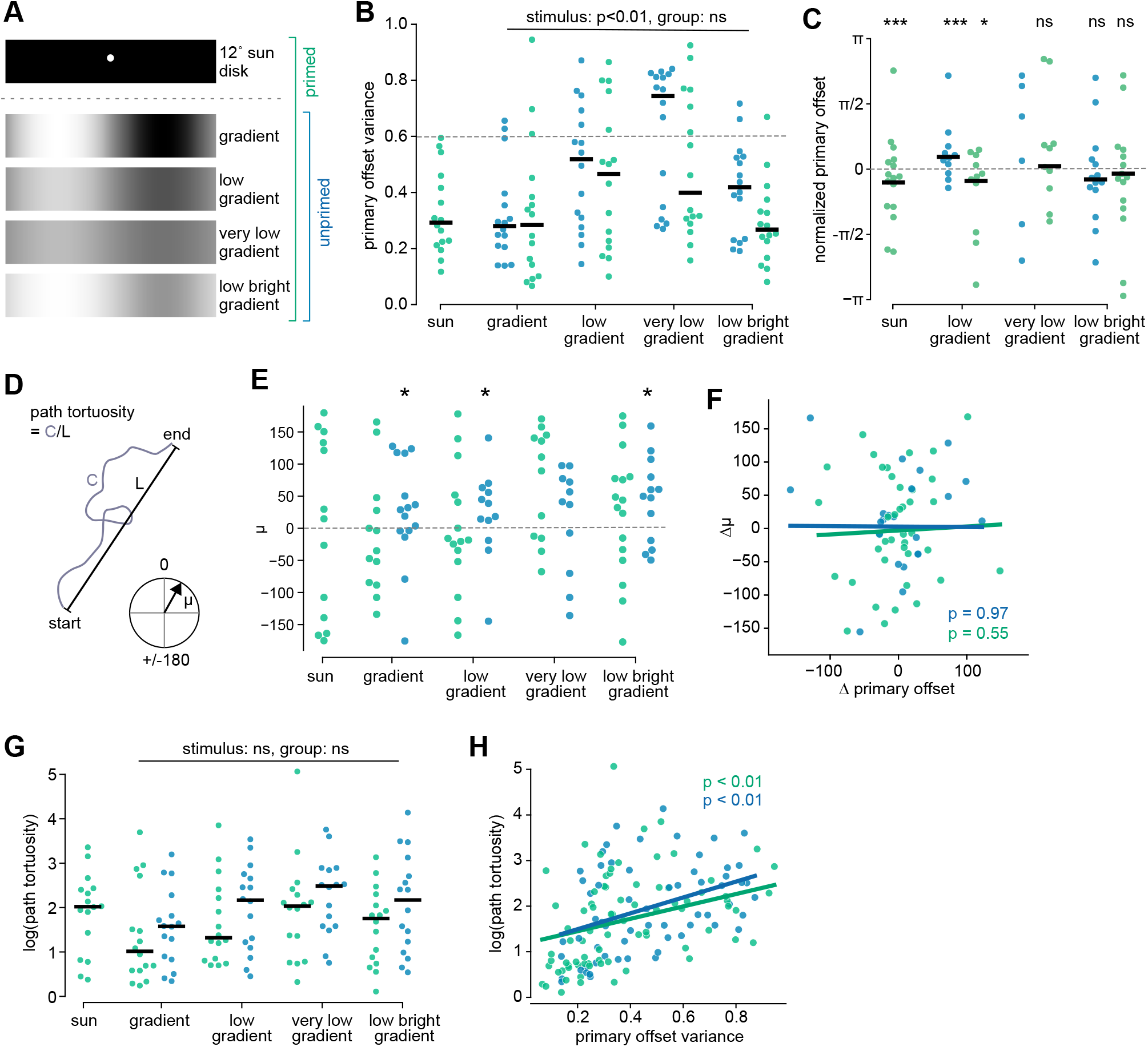
The effect of “priming” on compass stability. **A:** We used two experimental protocols. In one (primed, green, n=16), flies experienced a sun disk trial before trials with gradient stimuli. In the other (unprimed, blue, n=16), flies directly experienced the gradient stimuli. **B:** Distribution of the primary EPG offset variance across flies per trial, comparing the two experimental groups, primed (green) and unprimed (blue). Only datapoints where the primary offset was used for at least 50% of the trial were considered. Black bars mark median values per group and trial. For the gradient trials, we found that the visual stimulus had a strong effect on primary offset variance (p=0.0001, ANOVA) and the experimental groups had some effect (p=0.0782). We did not detect any significant interaction between the two variables (p=0.5566). See **Methods** for statistical details. **C:** Same as B, but for the normalized primary offset. Only datapoints where the primary offset variance was less than 0.6 are shown. For groups with at least 10 samples left after filtering, we show the circular mean and performed a Rayleigh test to test for deviation from a uniform circular distribution. Stars represent significant deviation from a circular uniform distribution (* p<0.05, ** p<0.01, *** p<0.001). See **Methods** for details. **D:** Illustration of the fixation direction and path tortuosity. **E:** Distribution of fixation direction across trials for primed and unprimed flies. Unprimed flies are more likely to perform negative phototaxis. Stars represent significant deviation from uniform circular distribution (*p<0.05 for a Rayleigh test). **F:** Correlation between changes in offset and changes in fixation, pooled across trials and flies. Trials where no fixation was detected and/or if the offset variance was above 0.6 were excluded (between 1 to 11 trials were excluded for a group, see **Methods**). Color code for groups as illustrated in A. *The grey line and p-value indicate the results of a linear mixed effects model with offset variance as the fixed effect and flies as random effects. **G:** Distribution of path tortuosity values across flies per trial, comparing primed and unprimed flies as in F. **H:** Correlation between path tortuosity and offset variance. Trials with an offset variance above 0.6 were excluded. Color code for groups as illustrated in A.

However, when we primed the fly with the sun disk environment, we observed that the stability of compass dynamics was enhanced even under challenging gradient conditions (**Figure 3B**). The stability of the compass was also improved by the brightness of the gradient (compare “low gradient” with “low bright gradient” stimuli in **Figure 3B**). The offset itself was in some cases preserved across all gradients we tested (**Figure 3—Figure Supplement 2A,C**). Across the population of tested flies, the offset was preserved between the sun, gradient and lower intensity gradient environments, but the distributions of normalized offsets were much wider at low contrasts (**Figure 3C**, note the unimodal circular distributions for sun and low gradient stimuli, indicating on average no change relative to the offset in the gradient environment). Thus, the fly compass does lose some stability in low intensity gradients, but the plasticity of the compass system allows prior exposure to a stronger gradient to ameliorate the situation somewhat, while also preserving the actual offset.

Thus far, we assessed the fly’s ability to perform menotaxis under sun disk and gradient conditions, and, separately, evaluated the stability of its compass. We now asked if the stability of the compass was perhaps more directly correlated with the fly’s ability to maintain a stable heading relative to its surroundings. Using the fact that our imaging experiments were performed in flies that were actively behaving on the ball, we quantified the flies’ behavior by their preferred heading direction within the visual environment, and by the straightness of their walking path (**Figure 3D**). We first looked for evidence of menotaxis across the different gradient environments (**Figure 3E**). While flies that were primed with an initial sun environment chose arbitrary walking directions, unprimed flies showed negative phototaxis, with their preferred walking directions clustered around the dark parts of the gradient (**Figure 3E**). Next, we looked for correlations between changes in the offset and changes in the fly’s heading preferences. Consistent with previous reports that have suggested that flies can change their heading preferences within a visual scene without changing the offsets of their compasses to that scene (Green et al., 2019), we found no correlation between changes in heading preference and changes in the fly’s offset (**Figure 3F**). Although changes in the offset may not directly relate to changes in flies’ heading preference, the compass is required for menotaxis. Thus, we next checked if stability in compass dynamics was correlated with flies maintaining straight paths in our imaging setup. We found that an increase in variability in the offset, our measure of the compass instability, was correlated with the tortuosity of flies’ walking paths (**Figure 3G,H**), and with a reduction in their net displacement during each trial, but not their absolute walking distance (**Figure 3 Supplement 2E,F**). Thus, we found that changes in the compass offset have no direct relationship with flies’ preferred headings, but compass stability likely affects the fly’s ability to keep a straight bearing.

Our results thus far suggested that either a sun disk or an intensity gradient in a virtual sky can serve as sufficient references for the stable operation of the fly’s internal compass. When presented with multiple compass cues simultaneously, many insects show a preference to follow information from one over the other (Dyer and Gould, 1983; el Jundi et al., 2014b; Wehner and Muller, 2006). To test if flies would show a preference for updating their compass system based on the sun disk or intensity gradient, we presented both stimuli simultaneously. In most natural daytime settings, the intensity gradient is brightest at the sun’s position in the sky. We compared the fly’s compass response between this naturalistic condition and a condition in which these cues—a localized bright light source and a widefield intensity gradient—are in conflict (**Figure 4A**), as is less likely in naturalistic settings, but can happen in twilight or under cloudy skies in the presence of artificial light sources. In both situations, we began by allowing the fly to experience a virtual sky with only the sun disk present, followed by a trial with both the sun disk and the intensity gradient and a trial with just a low intensity gradient, before returning the fly to the sun disk setting in a final trial. We found that the compass offset did not change dramatically when moving from a sun-like setting to settings that have intensity gradients if the brightest part of the gradient overlapped with the position of the sun disk (**Figure 4B**). However, when the two visual cues were placed in conflict, the compass either maintained its sun-tethered offset or switched to an offset 180° away from the original and consistent with tethering to the intensity gradient (**Figure 4C**). Curiously, in some cases, this altered offset was preserved when the fly was returned to the setting with just a sun disk (rightmost column of **Figure 4C**). We found that the stability of the compass did depend on the stimulus conditions (**Figure 4D**), but, surprisingly, not on whether the sun disk and gradient cues were presented in a naturalistic or conflicting manner. Furthermore, the increased compass stability in environments with both the sun and intensity gradient cues did not result in significantly straighter paths (**Figure 4E**), as seen previously with only the gradient stimuli (**Figure 3**). More in line with our expectations, individual flies that experienced the naturalistic sun disk and gradient stimulus showed more consistent heading preferences across trials (**Figure 4F**) than the group of flies that experienced the conflicting configuration of the sun disk and gradient (**Figure 4G**). Taken together, these findings suggest that sun and gradient cues may be represented in the fly brain by a shared population of neurons (see **Discussion**) and that a combination of the sun disk with the widefield intensity gradient increases the stability of the compass system. In summary, the fly’s compass tethered stably to the fly’s surroundings across the naturalistic variations in skylight that we probed in VR, and flies maintained menotactic preferences across these conditions to some degree.

**Figure 4:**
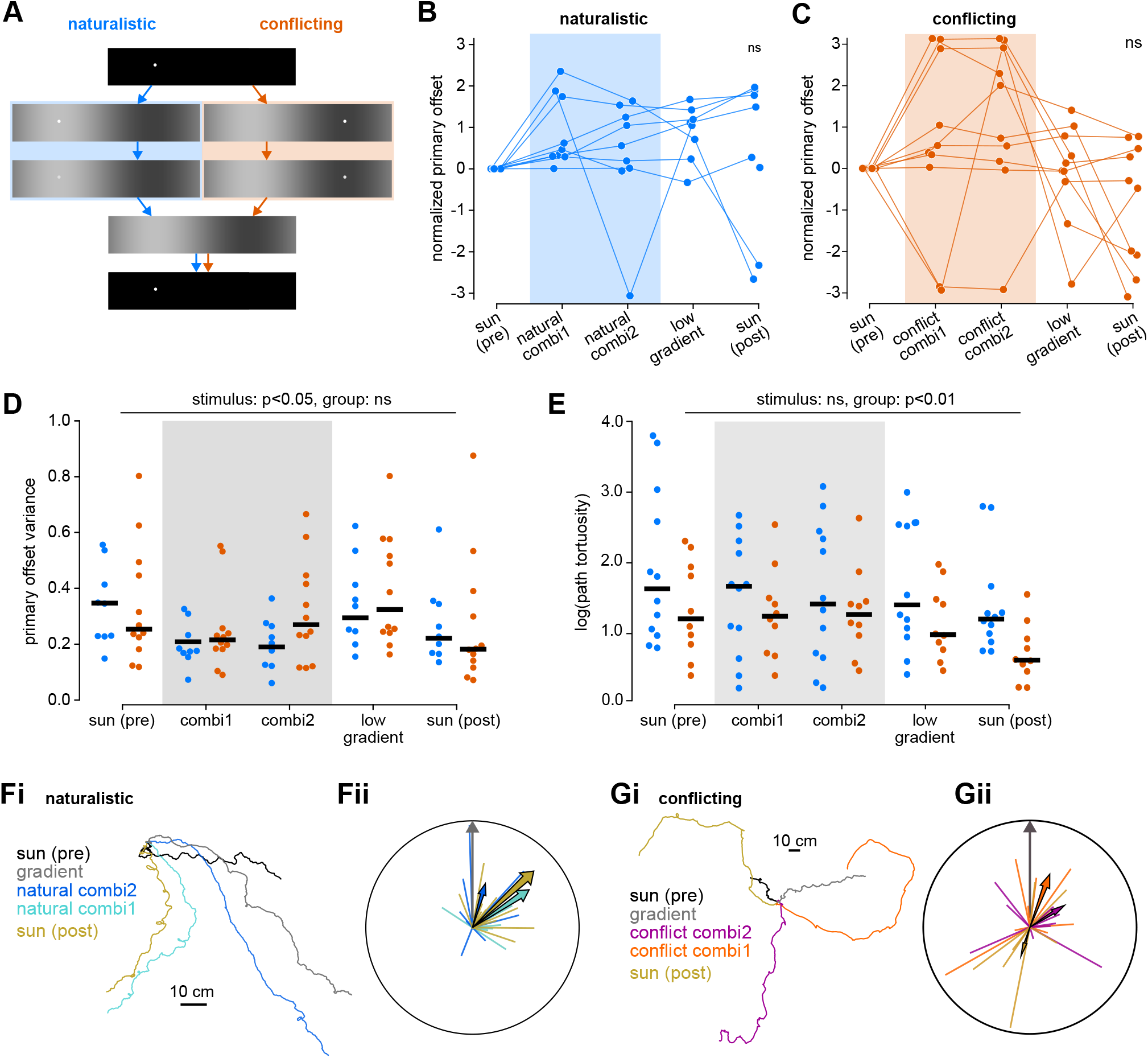
Naturalistic and conflicting compound stimuli affect compass stability and behavior differentially. **A:** Schematic of the experimental protocol. One group of flies experienced sun and gradient in a naturalistic configuration (sun on bright part of gradient, n=12, blue), the other group in a conflicting configuration (sun on dark part of gradient, n=12, orange). All female SS00096 > GCaMP7. **B:** Normalized offset location for the naturalistic stimulus group. The offset location during the first sun trial was used as the reference. Only trials where the circular variance was below 0.6 and the primary offset was used for at least 50% of the trial time are shown (see **Methods** and **Figure 2 Supplement 1**). **C:** Same as B but for the conflicting stimulus group. **D:** Comparison of the circular variance in the naturalistic and conflicting stimulus group. Only trials where the primary offset was used for at least 50% of the trial time are shown. The black bars indicate the median. An ANOVA (see Statistics for details) found that the visual stimulus (p=0.0301) but not the experimental group (p=0.3502) contributed to explaining the variance in the sample, nor was there an interaction between group and stimulus (p=0.4740). **E:** Comparison of the path tortuosity in the naturalistic and conflicting stimulus group. ANOVA showed that the experimental group contributed to explaining the variance in the sample (p<0.01) but the stimulus had no effect. **F:** Fixation directions of flies in the naturalistic stimulus group. **Fi:** Walking paths of a single fly taken across the five trials. **Fii:** Fixation plot as explained in **Figure 1G** for fixation directions (normalized to the fixation direction in the low gradient) chosen by flies in the naturalistic stimulus group. **G:** Same as F but for the conflicting stimulus group.

Having established the performance of the fly compass in a variety of different sky-like visual VR conditions, our final set of experiments challenged the fly with still more naturalistic surroundings, settings with local cues in the form of virtual grass patches set against either the intensity gradient or the sun disk (**Figure 5A,B**). Inspired by the observation that insects rely on UV and blue light to detect celestial cues, we chose to keep the skylight cues bright while making terrestrial grass objects dark, as plants absorb UV light (Barta and Horvath, 2004; Moller, 2002). To do this, we chose a sky with a bright sun disk on a grey background and the intermediate low gradient (**Figure 1—Figure Supplement 2**). The sun disk on a grey sky is a weaker compass cue compared to sun disk on a black sky (**Figure 5—Figure Supplement 1**), but the compass stability of this condition is comparable to that with a strong intensity gradient (compare to **Figure 3B**). For both skylight conditions, we tested different grass patches densities (**Figure 5A,B**). The fly was allowed to walk up and through the patches, which meant that the sun disk or the brightest part of the gradient could sometimes be obscured by the grass, as might occur in natural foliage. We expected that with increasing occlusion of celestial cues, the compass would become unstable. Indeed, we found that when the grass was densest and flies’ only global cue was a sun disk, which would often be completely obscured, compass dynamics were unstable (**Figure 5C,G**). By contrast, many flies in the gradient environment were able to maintain a stable compass even in the dense grass environment, likely because the wider intensity gradient was still sufficiently visible (**Figure 5C,H,I,** and **Figure 5 Supplement 2A,B**). We expected that the high-contrast terrestrial objects paired with distal compass cues would create conflicts for the compass, because of the variety of dark and light patches at different orientations that seem likely to trigger responses from visual ring neurons (Hardcastle et al., 2021; Omoto et al., 2017; Seelig and Jayaraman, 2013; Shiozaki and Kazama, 2017; Sun et al., 2017). Such conflicts, paired with the partial or—in the case of the sun disk—complete occlusion of the celestial cues, may have resulted in offset changes across trials (**Figure 5D,E**). Evidence for conflicts between local objects and sky cues destabilizing the compass system also came in the form of transient offset instabilities we observed when flies transitioned through grass patches (**Figure 5H,I**). Especially in the sparse grass condition with an intensity gradient sky, some flies showed multiple, locally stable offsets (**Figure 5F, J**). Overall, we concluded that if a sufficiently large area of the sky is visible to the fly, the visual inputs to the fly’s compass provide it with the necessary stability to tether to global cues, allowing it to remain stable through periods when these cues might either be compromised or transiently obscured. This presumably allows the fly to maintain its pursuit of behavioral goals that rely on this internal representation even in the presence of visual clutter.

**Figure 5:**
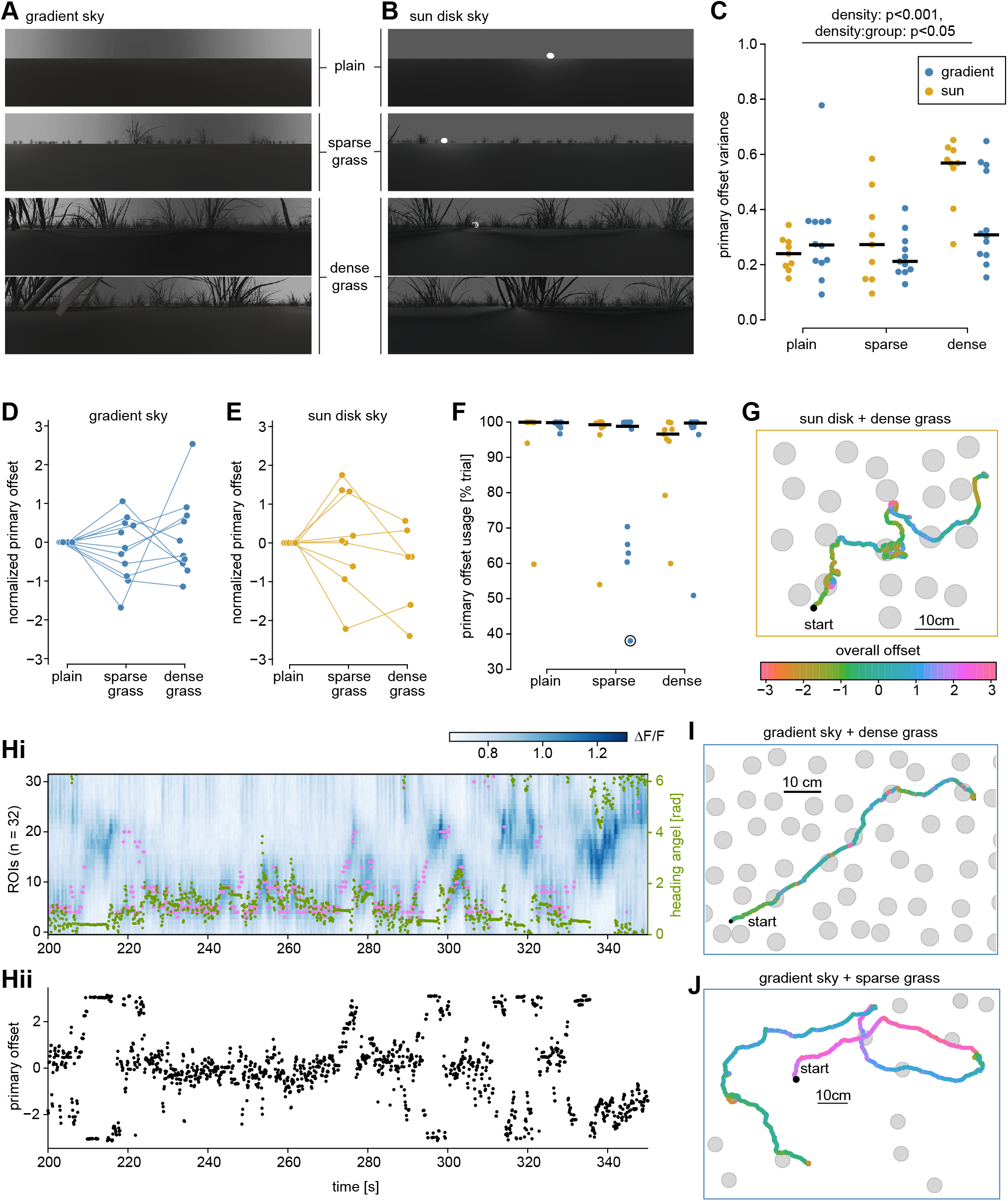
Compass stability in cluttered environments. **A:** Panoramic views of the grass worlds with an intensity gradient sky. Three densities were tested, one plain world with no grass, a sparsely and a densely populated world. **B:** The same as A, but for the grass worlds with a sun disk sky. **C:** Primary offset variance across different grass densities comparing the group with a sun disk background (yellow, n=9) and gradient background (blue, n=12). Black bars mark the medians. Increasing the density of grass objects significantly increases the offset variance (p=0.0001, ***, ANOVA effect of density). Further ANOVA indicates there is an interaction between the density and the sky cue (p=0.03009, *). **D:** Normalized primary offset across grass density in the gradient sky group. All offsets were normalized to the respective value in the world without grass (plain). **E:** Same as D, but for the group with a sun sky. **F:** Percent of the trial time, during which the primary offset was used. For most flies the primary offset is used for nearly the full trial time. However, for the gradient sky a subset of flies shows uses a second offset for a large amount of the trial time (primary offset only around 60%), and in the sun sky group the offset usage drops slightly in the dense grass world. The circled dot corresponds to the fly shown in J. **G:** An example walking trajectory of a fly in the dense grass world with a sun sky. The grey circles indicate the position of grass tussocks (compare to B). The starting point is indicated by a black dot. The trajectory is color-coded by the offset location (not the primary, but the overall offset that is defined for all time points, see **Figure 2** and **Methods**). **Hi:** Calcium activity in EPG neurons in the EB as shown in **Figure 2D** for a fly in the dense grass environment with a gradient sky. The peak position (pink) and the head direction (green) are overlaid. **Hii:** The primary offset for the calcium activity shown in H. **I:** The walking trajectory of the same fly as shown in H, visualized as described in G. **J:** Walking trajectory of a fly in the sparse grass environment with a gradient sky, visualized as described in G. Note the changes between multiple stable offsets.

## DISCUSSION

We found that the fly’s neural compass is stable in the presence of skylight cues such as sun-like disks and intensity gradients, and that it can maintain its spatial relationship to these global cues even in the presence of local landmarks. Our results suggest that sun-like objects and intensity gradients akin to those they might produce in the sky are both effective in updating the fly’s neural compass, and either is sufficient for goal-directed navigation in form of menotaxis (**Figure 1-3**). However, these cues can offer distinct advantages in terms of supporting a stable head direction estimate. When flies navigate in challenging intensity gradients that span a limited contrast range, prior exposure to a sun disk significantly improved compass stability as well as the fly’s ability to walk straight during menotaxis, emphasizing the role of plasticity in the compass system (**Figure 3**). On the other hand, in environments where local objects could occlude distal skylight cues, the compass system remained more stable in the presence of widefield intensity gradients than with the sun disk (**Figure 5**). As might be expected for a system that has evolved to exploit available cues in nature, a combination of the sun disk and intensity gradient stabilized the compass system better than conditions in which only one of the two cues was available (**Figure 4**).

The ability of flies to keep a stable offset—that is, to maintain a common frame of reference for orientation—across visually distinct settings is likely the result of specialized processing within the anterior visual pathway (AVP) (Hanesch et al., 1989a; Hulse et al., 2021; Omoto et al., 2017; Timaeus et al., 2020) that provides visual information to the compass. A simple mechanism for generating consistent offsets when transitioning between a sun and intensity gradient environment would be that both visual stimuli appear similar to the compass system. Indeed, it has been suggested that deriving orientation information does not require fine visual feature discrimination (Dewar et al., 2015; Dewar et al., 2017; Wystrach et al., 2016). In fact, the relatively large size of ring neuron receptive fields (Hardcastle et al., 2021; Omoto et al., 2017; Seelig and Jayaraman, 2013; Shiozaki and Kazama, 2017; Sun et al., 2017) might act like spatial filters and ensure that these neurons respond similarly enough to sun disk cues and larger-scale light gradients to evoke consistent compass neuron responses whether the sun is visible or hidden behind clouds..

In conditions where multiple bright features are present, such as when we presented a sun disk and intensity gradient in a conflicting configuration, our results suggest that one of the features is selected to tether the compass (**Figure 4C**). This is consistent with proposed stimulus selection mechanisms in input circuits to the compass (Hulse et al., 2021). Inhibitory visual ring neurons at the last stage of the AVP likely inhibit each other at their outputs in the EB (Hulse et al., 2021; Isaacman-Beck et al., 2020). This potential mechanism for stimulus selection (Hulse et al., 2021) could preferentially bias the compass to receive information from ring neurons responsive to a dominant visual feature. In the conflict condition we tested it is possible that no one feature was clearly dominant, resulting in the bimodal split across the population of measured flies. Our results, including a test of different sun disk sizes and contrasts (**Figure 5—Figure Supplement 1**), suggest that the system exhibits some invariance to feature size but is sensitive to local contrast. What exactly determines dominance in a visual feature—for example the size or intensity—has yet to be determined for this circuit.

Finally, plasticity in the mapping between visual features and compass neurons (Cope et al., 2017; Fisher et al., 2019; Green and Maimon, 2018; Kim et al., 2019) provides a mechanism for ensuring that the compass system is more stable with increased exposure to visual features with a stable angular relationship to each other and movement across the fly’s field of view that is consistent with the fly’s turning behavior, both of which would favor global cues over local ones (Kim et al., 2019). This could prevent tethering of the compass to blades of grass that the fly walks by in our VR. We note that our virtual skies included neither spectral nor polarized light information—skylight cues that insects are likely to rely on (Brines and Gould, 1979; el Jundi et al., 2015; Homberg, 2004; Warren et al., 2018; Wehner, 1997; Weir and Dickinson, 2012), and which are known to be present in the AVP (el Jundi et al., 2014a; Hardcastle et al., 2021; Heinze and Reppert, 2011; Vitzthum et al., 2002). These cues are likely to further stabilize the compass system in cluttered settings and transiently changing skylight conditions.

Studies across many insects have established the importance of celestial cues not only for updating an internal head direction representation but also directly for navigation (Dacke et al., 2014; el Jundi et al., 2014b; Kraft et al., 2011; Mathejczyk and Wernet, 2019; Warren et al., 2018; Wehner, 1997; Weir and Dickinson, 2012). Recent work in flies has used silencing experiments to link menotaxis in that species to the compass system (Giraldo et al., 2018; Green et al., 2019; Turner-Evans et al., 2021). Consistent with such results, we found that silencing the compass neurons led flies to perform negative phototaxis. We note that this behavior is different from the well-known frontal fixation response of walking flies to dark stripes (Bahl et al., 2013; Horn, 1978; Horn and Wehner, 1975; Strauss and Pichler, 1998). Whether this behavior may be dependent on the contrast conditions—dark-on-bright versus bright-on-dark cues—is unclear. In general we found that the preferred arbitrary walking direction (goal direction) of individual flies across different environments that we evaluated over the time scales of minutes appeared unrelated to changes in the offset (Green et al., 2019). Instead, we found that even in flies with an intact compass system the average offset stability was, under certain conditions, correlated with prevalence of negative phototaxis instead of menotaxis. Furthermore, the compass stability was correlated with the straightness of walking in flies during menotaxis.

An age-old dilemma in neuroscience concerns the challenges and value of using natural stimuli in probing neural circuits and animal behavior. In this study, we provide a powerful framework to study the fly compass system during compass-guided navigation in naturalistic settings, which enables direct and simultaneous comparisons of neural activity and behavior. We hope that this will provide opportunities to bridge between the level of circuit understanding that can be achieved with well-controlled stimuli and the ethological insights that have been gleaned from studying animals in their natural habitat.

## ACKNOWLEDGMENTS

We thank Frank Loesche and Michael Reiser for advice on using fictrac for ball tracking, Pratyush Kandimalla for helpful discussions about ring neurons and feedback on the manuscript, and Rebekah Zhang for help with converting an existing VR rig for behavior to the newer Unity-based system. We are grateful to Vasily Goncharov for assistance with the two-photon microscope and Davis Bennett for useful discussions on rig design. We thank Dan Turner-Evans, Brad Hulse for advice over the course of the study, and for feedback on the manuscript.

SC was supported by the Janelia Undergraduate Scholar program during the pilot phase of the project. This work was supported by the Howard Hughes Medical Institute.

## AUTHOR CONTRIBUTIONS

HH, AMH and VJ conceived the study. HH and TG designed and constructed the projector-based display for the imaging rig. PH developed software for the virtual reality system with input from HH. HH assembled the VR rigs used in imaging and behavior experiments. HH and SC performed calibrations and experiments. HH and SC analyzed the data with input from VJ and AH. HH and VJ wrote the manuscript with input from SC.

## METHODS

### Fly strains

To target EPG neurons in calcium imaging experiments, we used the SS00096 split Gal4 line. We crossed SS00096 to 20XUAS-IVS-jGCaMP7f (Dana et al., 2019), resulting in GCaMP7f expression in EPG neurons. We used flies of this genotype for all imaging and most behavior experiments. We also used SS00096 to silence EPG neurons in behavior experiments, by crossing this line to WTB;;UAS-Kir2.1.eGFP (generated by Scarlett Coffman, FlyCore, Janelia Research Campus, Ashburn, VA, USA and gifted to us by Chuntao Dan). WTB stands for wild type Berlin. To generate a control genotype for the silencing experiments, we crossed SS00096 to WTB.

### Fly rearing

Flies were reared in small fly vials with cornmeal media in incubators at 23°C with 60% relative humidity and a 16:8 h light:dark cycle. To prepare 10 l of corn meal food, 7.23 l of water and 59.66 g agar (fly agar, Tic Gums Inc, Belcamp, MD, USA) were brought to a boil. 160.68 g yeast (inactive dry yeast, Genesee Scientific, San Diego, CA, USA) and 664.84 g cornmeal (Quaker Yellow Corn Meal, Quaker Oats Company, Chicago, IL, USA) were mixed with 1.59 l of water. The yeast/cornmeal mixture was added to the boiling agar solution. 0.4 l molasses was then added and the mixture was simmered. Finally, after turning off the heat, 42 ml of Propionic acid and an antifungal agent, Tegosept (79.55 ml, Genesee Scientific, San Diego, CA, USA), were added. To limit the density of animals within a food vial and to ensure good food conditions, crosses were flipped every 4-5 days and offspring were transferred to fresh vials every 2-3 days. The latter enabled control of the age of experimental flies.

### Preparation of flies for experiments

For both behavior and imaging experiments, we used 4-8 d old female flies. To encourage walking activity, flies were wet starved for 22-26 h before the experiment by transferring flies into vials with a moist filter paper but no food. This step was carried out without cold anesthesia.

### Behavior experiments

An hour before the experiment, the flies were transferred into a tube and cold anesthetized at ~4°C. They were then transferred onto a cold plate and sex-sorted. A thin tungsten wire (tether) was glued to the back of a fly by applying UV-curable glue on the tip of the wire, lowering the tip onto the fly’s thorax, and shining UV light for 10-20s. The flies were head-fixed to reduce head movements that interfere with closed-loop visual stimulation. This was done by adding a drop of glue to the neck connective and curing it with UV light. The flies were allowed to recover from the cold anesthesia for at least 15 minutes before starting the experiment. The tether was inserted into a custom holder and gently lowered onto the spherical treadmill using micromanipulators. This allowed for precise positioning of the fly on the ball.

### Imaging experiments

Starved flies were transferred to an empty vial, cold-anesthetized for less than a minute at ~4°C and transferred onto a cold plate. The proboscis was pushed into the head capsule and fixed with a small amount of wax. Because some flies tended to not be as active after this step, we prepared several flies, let them recover from anesthesia and selected individuals for experiments that showed high levels of activity. Next, a fly was glued to the tip of a thin tungsten wire (a tether) with UV-curable glue (KOA 300, KEMXERT, York, PA, USA). Gluing the fly to this tether enabled precise positioning under a custom-designed shim. The shim consisted of a pyramid made from a folded metal sheet glued into a frame that could hold saline (**Figure 1C** inset). The pyramid had a small hole at the tip, into which the fly’s head was placed, covering the upper part of the eyes. The shim was designed to provide a field of view of at least 20° above the horizon to the fly (previous models ranged from 9-12°). Using a viscous UV-glue (Fotoplast Gel, Dreve Dentamid GmbH, Unna, Germany) the fly’s head was glued to the pyramid, closing the hole. Saline was then added and a small hole cut into the cuticle on the top of the fly’s head. The cuticle and fat were removed, and air sacs were pulled back, providing a clear view of the central brain. The saline had the following composition (in mM): NaCl (103), KCl (3), TES (5), trehalose 2 H_2_O (8), glucose (10), NaHCO_3_ (26), NaH_2_PO_4_ (1), CaCl_2_ 2 H_2_O (2.5), MgCl_2_ × 6 H_2_O (4), dissolved in Milli-Q filtered water.

### Calcium imaging in behaving flies

For calcium imaging experiments, we used a custom-built two-photon microscope (Kim et al., 2019) running ScanImage (version 2018b (Pologruto et al., 2003), Vidrio Technologies). For the excitation, we used a tunable Ti:Sapphire laser (Chameleon Ultra I, Coherent) with the wavelength set to 930 nm and ~8 mW power at the sample (scanning mode). To image the EB, we acquired image volumes consisting of 8 planes (150 x 150 pixels each), spaced 6 μm apart with two additional discard planes. This resulted in a volume acquisition rate of 9.55 Hz. We used a 40x objective (Nikon CFI APO NIR Objective, 0.80 NA, 3.5 mm WD), whose long working distance helped to minimize occlusions from the fly’s field of view. To free up space around the objective for the panoramic screen, we flipped the orientation of the PIFOC objective scanner (Physik Instrumente, Auburn, MA, USA) by 180°. The microscope head and objective were tilted by 30° to reduce the number of planes necessary for covering the complete EB with the fly’s head at a natural angle (**Figure 1C** inset). Before calcium imaging experiments, flies were positioned on the treadmill ball and given 10-20 minutes to acclimate. During this time the acquisition and virtual reality software were initialized, and imaging volume was selected. The experiment was aborted if the signal was low or if there was too much brain movement. The temperature around the fly was ~25 °C and the humidity was ~30%.

### VR setup

We used two VR setups for this study: one for behavior experiments in non-dissected flies (modified from (Haberkern et al., 2019)), and a second one for imaging experiments (**Figure 1C**, **Figure 1 Supplement 1**). Both setups consisted of a spherical treadmill and a panoramic screen, and used the same Unity-based VR software (see below)

### Spherical treadmill and ball tracking

The ball holder for the spherical treadmill has been described in detail in (Seelig et al., 2010). The ball was machine-cut from polyurethane foam (FR-7110, Last-A-Foam, General Plastics Manufacturing Company, Tacoma, WA, USA), weighted about 50 mg, and had a diameter of 8.8 mm in the behavior rig, and 9.1 mm in the imaging rig. To allow for low-friction rotation of the ball, it rested on an air cushion maintained by a constant air flow (0.55 l/min, controlled with a mass flow controller by Alicat Scientific, Tucson, AZ, USA). The ball motion was captured by a high-speed camera in infrared illumination (Grasshopper3 GS3-U3-23S6M, 2.3 MP, Teledyne FLIR, Wilsonville, OR, USA). The frame rates for the ball tracking were 165 Hz on the behavior rig and 146 Hz for the imaging rig. We used FicTrac (Moore et al., 2014) to process the video data to estimate the orientation of the ball and reconstruct the virtual trajectory of the fly. The output from FicTrac was sent to a socket port, where it could be read out by the VR software. Tracking with FicTrac requires the ball surface to be patterned such that there is a unique pattern for each orientation. We painted high-contrast patterns on the ball with black acrylic paint on a red base coat (Premiere Acrylic Colour, Laurel, NJ, USA).

### Visual display

In both setups, we used DLP projectors to back-project images onto a panoramic screen, which was made from a white diffuser sheet (V-H105-CV07 for the imaging rig and V-HHDE-PM06-S01-D01 for the behavior rig, both from BrightView Technologies, Durham, NC, USA). The screen geometry and projectors differed between the two setups (see below for details and **Figure 1—Figure Supplement 1**), but both setups used the same external illumination source (SugarCUBE LED Illuminator, Edmund Optics Inc, Barrington, NJ, USA), which was attached to the customized projectors. The light was blue with a peak at 458 nm wavelength, resulting in monochrome blue images for the VR stimulus. Blue light was chosen to simplify filtering out light-noise in two-photon imaging experiments.

### Behavior rig

The hardware for the behavior setup has been described in detail previously (Haberkern et al., 2019) and dimensions are shown for reference in **Figure 1—Figure Supplement 1C,D**. The rig used two projectors, each generating an image with 720 x 1280 pixel resolution at 120 Hz refresh rate for a continuous panoramic 1440 x 1280 pixel display. We did not use frame packing as described before (Haberkern et al., 2019), and thus, the frame rate was equal to the refresh rate. The fly was placed symmetrically between the two faces of the V-shaped display, 3.8 cm from the center of the display and at 0.28x the screen height (see **Figure 1—Figure Supplement 1C,D**). The screens covered 115° of the fly’s azimuthal field of view on both sides (230° azimuthal coverage in total). The distance of the screens from the fly varied along the azimuth (2.7 cm to 7.8 cm), resulting in variations of the display coverage in elevation (77°to 58°). The view below the horizon was restricted to −50° by the ball surface.

### Imaging rig

The visual display in the imaging rig was formed by a pentagonal panoramic screen. For a fly positioned in the center of this display, the screens spanned a field of view of 144° azimuth to both sides (288° azimuthal coverage in total) and 76°-92° in elevation (see **Figure 1 Supplement 1A, B** for details). The custom-build frame was built for the screen using five 6 mm posts (MS3R, Thorlabs) and custom 3D-printed braces. The visual stimulus was rear projected onto the four screens using four projectors (DLPDLCR2010EVM, DLP® LightCrafter™ Display 2010 Evaluation Module, Texas Instruments). Each projected image had a resolution of 480 x 720 pixels, together creating a 1920 x 720 pixel panorama. The framerate of the projectors, as well as the monitor connected to the setup computer, was set to 144 Hz. The projectors were modified for close-range projection to accommodate the setup in the confined space around the microscope. Further, the built-in LEDs were disconnected, and a light guide was fitted onto the projector’s optical module. An external blue light source was connected to each projector via the light guide. To limit noise from the projector light during imaging experiments, an additional shortpass filter with 500nm cutoff was inserted into between the light source and the light guide (Everix Ultra-Thin Shortpass Filters Stock #35-892, Edmund Optics, USA).

### VR software

Building on the Unity game engine, we developed a novel, versatile VR software. We developed packages that extend Unity functionality in a modular manner to accommodate specialized needs for animal VR. These packages are published as part of the Janelia Unity Toolkit, and allow precise control of frame rate, setup of a panoramic display, a connection with FicTrac via socket connection, custom collision handling, data logging and more. Briefly, we used the Unity editor to design virtual worlds, largely in an automated fashion using imported packages from the Janelia Unity Toolkit. For instance, a customized perspective projection matching the display geometry of the respective rig was set up using the “Setup Cameras N-gon” and “Camera Utilities” packages. The “Force Render Rate” package was used to set and force a fixed frame rate, matching the chosen frame rate of the respective rig (120 Hz for the behavior rig, 144 Hz for the imaging rig). Several other packages implement a socket connection to read FicTrack ball tracking data, implement a custom positional updater based on the received data, and perform custom collision handling. The “Logging” package implements a data logging framework for writing an event-based log to a json file, and the “NIdaqMx” package implements an interface for communication with a NIDaq board (National Instruments, 781005-01 NI USB-6218 BNC BUS-POWERED M SE). Documentation for these and other packages can be found on github (https://github.com/JaneliaSciComp/janelia-unity-toolkit).

### Brightness correction for projected images

We found that the projectors applied a transformation to projected images, which meant that the output grayscale brightness did not scale linearly with the input pixel values. This transformation is often referred to as the “gamma curve” of the display. For both rigs, we obtained the gamma curve empirically by measuring the output brightness of the projectors with a power meter (PM100D with S130C sensor, Thorlabs) for equally spaced pixel values (**Figure 1—Figure Supplement 4,** note that the curves for different projectors of the same model did not differ). We then inverted the measured curves to obtain a projector-specific correction function. This function was applied pixel-wise to rescale texture images that were used in our VR environment, such that the projected images had the intended brightness distribution and contrast. We confirmed that these transformed textures appeared as intended on the display using intensity measurements.

### Generation of virtual environments

#### Distant panoramas

To simulate celestial compass cues, we generated virtual worlds in which a fly could control its orientation relative to a cylindrical panorama, but where translation had no effect on the visual environment. This effectively simulates visual panoramas that are infinitely far away from the fly. To generate these environments, a panoramic image that spanned 360° azimuth was rendered in Python (version 3.8, https://github.com/hjmh/vrStim) and then wrapped around a virtual cylinder in a Unity world using the “Background” package from the Janelia Unity Toolkit. The image was wrapped in an anti-clockwise manner around the virtual cylinder, starting at 90° to the right (**Figure 1—Figure Supplement 3B)**. The virtual cylinder was centered on the fly, so that the horizontal midline of the panorama was displayed at the fly’s horizon. The cylinder’s x/y-position was locked with respect to the fly’s position, ensuring that changes in the fly’s position had no effect on the displayed stimulus. An overview of the panorama images used in this study is given in **Figure 1—Figure Supplement 2**.

#### Immersive, two-dimensional worlds

The 2D immersive worlds were composed of the following components: a distant, unapproachable panorama, a textured ground plane, and, for some experiments, approachable objects. The **distant panorama** was created through a custom skybox. We generated the skybox textures from panorama images used in the “Distant panorama” worlds using Blender (version 3.0.0) and a custom python script for Blender that allows rendering of spherical images or movies (https://github.com/JaneliaSciComp/blender-spherical-video) from a 3D scene. We used two types of skyboxes, one with a low gradient (C2), and one with a white on grey sun disk (B2s). The textured **ground plane** was generated by creating a plane object positioned 1 mm below the camera (virtual fly) position. A greyscale texture image consisting of white noise (shown in **Figure 1—Figure Supplement 3B**) was mapped onto the plane object to ensure that the fly could see an optic flow stimulus when running across the plane. Finally, for some experiments, we added **approachable objects**, plant models, to the virtual world. These plant models were constructed from assets purchased on CGTrader and modified to be correctly loaded and displayed as Unity assets with different levels of detail to improve rendering performance. A Janelia Unity Toolkit package (“Meadow”) was used to programmatically place plant models in a stochastically jittered grid layout on the ground plane. The plant model, grid size, placement jitter and placement probability were specified in a JSON parameter file. We used the following conditions in our experiments:

- Plant model: “Grass clump”
- Grid space: A total grid space of 200.0 x 200.0 m large square
- Density and jitter: 20 x 20 grid cells, resulting in 10.0 x 10.0 cm large squares. Plant models were placed with a 3.33 cm jitter from the center of the cell.
- Placement density: For the *sparse grass* world, each grid cell was occupied with a grass with p=0.2, in the *dense grass* world with p=0.99.

We used a script to measure how much of the sky was visible when moving through the worlds with plants. We found that without any plant models, 26% of the displayed image showed the sky. In the *sparse grass* world 17-23% (often 20-22%) of the sky was visible, while in the *dense grass* it was only 8-20% (often 12-15%).

The **lighting** in immersive worlds was adjusted to be white light, and the direction of the directional light was adjusted to match the skybox image (15° elevation or x-rotation and 90° azimuth or y-rotation).

### Experimental protocols

Each experiment consisted of several trials (see below), which were each 8 min long. An overview of experiments, genotypes used, and number of samples collected is given in **Supplementary Table 1**. The experimental protocols are described below. To avoid systematic biases from the order of presentation, we varied the presentation order of different trials as described below.

#### Distant panorama experiments

##### Gradient transitions

Flies were presented with four types of gradient panoramas, which differed in the modulation depth and mean brightness (see **Figure 1—Figure Supplement 2B** for details): gradient (C1), low gradient (C2), very low gradient (C3), and low bright gradient (C4). Each fly was tested in each panorama environment for 8 min trials each. We tested two presentation orders alternatingly, C1 →C2 →C3 →C4 and C4 →C3 →C2 →C1.

##### Gradient transitions with initial sun

The *gradient transitions* assay was repeated, but each experiment began with an initial presentation of a sun disk panorama with a 12° sun disk (A2), resulting in the following two orders being tested: A2→C1→C2→C3→C4 and A2→C4→C3→C2→C1. The sun disk position was just above the horizon line.

##### Sun disk size test

We tested 5 different sun disk stimuli that varied in the size of the sun disk or the contrast to the background (**Figure —Figure Supplement 2A**). All sun disks were positioned just above the horizon line. With maximum contrast (white sun disk on black background), we tested 12° (A2), 6° (A2s), 3° (A2s3) and 1° (A2ss) large sun disks. For the 6° large disk, we also tested a bright on grey stimulus (B2s). We tested these 5 panoramas in 8 different orders (see **Supplementary Table 1** for details).

##### Combination of sun disk and gradient

We tested combinations between the 6° sun disk (A2s) and the low gradient (C2) panoramas that varied in the sun disk intensity and in the placement of the sun disk on top of the gradient panorama (**Figure 1—Figure Supplement 2C**). We ran two assays with these combination stimuli:

- <u>Naturalistic configuration</u>: In this assay combined sun disk and gradient panoramas had the sun disk positioned in the center of the bright peak of the gradient, in one case the sun was white (C2S1s), in the other case it was light grey (C2S2s). We tested these stimuli together with the low gradient without a sun disk (C2) as well as initial and final presentations of the high contrast 6° sun disk (A2s). We looked at two presentation orders: A2s→C2S1s→C2S2s→C2→A2s (disappearing sun), A2s→C2→C2S2s→C2S1s→A2s (appearing sun).
- <u>Conflicting configuration</u>: The second assay followed the same logic as the assay with naturalistic configuration, but here the panoramas with a sun disk and gradient had the sun disk positioned on the dark peak of the gradient profile (C2T1s, C2T2s). We tested two orders here as well: A2s→C2T1s→C2T2s→C2→A2s (disappearing sun), A2s→C2→C2T2s→C2T1s→A2s (appearing sun).

#### Experiments with immersive, 2D VR worlds

We used six different 2D virtual environments that differed in local object density and in the presented distal cues. See above in “Generation of virtual environments” for details on how these worlds were constructed. We ran two assays, where we tested three different densities of local grass objects (no grass, sparse grass, and dense grass), but kept the distal cue constant. In one assay we tested a low gradient skybox (Gradient with grass), in the other we used a sun disk skybox (Sun disk with grass). See **Figure 5** for an illustration for these 2D worlds. For both assays we varied the presentation order as detailed in **Supplementary Table 1**.

**Supplementary Table 1:** Overview of experiments.

| Assay | Rig | Genotype | Orders | n | N |
| --- | --- | --- | --- | --- | --- |
| Gradient transitions | behavior | SS96 x GCaMP7f | C1 C2 C3 C4 | 20 | 40 |
|  |  |  | C4 C3 C2 C1 | 20 |  |
|  | imaging | SS96 x GCaMP7f | C1 C2 C3 C4 | 8 | **16** |
|  |  |  | C4 C3 C2 C1 | 8 |  |
| **Gradient transitions primed with sun** | behavior | SS96 x GCaMP7f | A2s C1 C2 C3 C4 | 20 | **20** |
|  |  | SS96 x Kir | A2s C1 C2 C3 C4 | 20 | **20** |
|  |  | SS96 x WTB | A2s C1 C2 C3 C5 | 20 | **20** |
|  | imaging | SS96 x GCaMP7f | A2 C1 C2 C3 C4 | 8 | **16** |
|  |  |  | A2 C4 C3 C2 C1 | 8 |  |
| **Naturalistic gradient sun disk combination** | behavior | SS96 x GCaMP7f | A2s C2S1s C2S2s C2 A2s | 10 | **20** |
|  |  |  | A2s C2 C2S2s C2S1s A2s | 10 |  |
|  | imaging | SS96 x GCaMP7f | A2s C2S1s C2S2s C2 A2s | 6 | **12** |
|  |  |  | A2s C2 C2S2s C2S1s A2s | 6 |  |
| **Conflicting gradient sun disk combination** | behavior | SS96 x GCaMP7f | A2s C2S1s C2S2s C2 A2s | 10 | **20** |
|  |  |  | A2s C2 C2S2s C2S1s A2s | 10 |  |
|  | imaging | SS96 x GCaMP7f | A2s C2S1s C2S2s C2 A2s | 6 | **12** |
|  |  |  | A2s C2 C2S2s C2S1s A2s | 6 |  |
| **Sun size test** | imaging | SS96 x GCaMP7f | A2 A2s A2s3 A2ss B2s | 2 | **16** |
|  |  |  | A2s A2 B2s A2ss A2s3 | 2 |  |
|  |  |  | B2s A2s3 A2 A2s A2ss | 2 |  |
|  |  |  | A2ss B2s A2s A2s3 A2 | 2 |  |
|  |  |  | A2s3 A2ss B2s A2 A2s | 2 |  |
|  |  |  | A2s A2ss A2s3 B2s A2 | 2 |  |
|  |  |  | A2 A2s3 B2s A2s A2ss | 2 |  |
|  |  |  | A2s3 B2s A2ss A2 A2s | 2 |  |
| **Gradient comparison (pilot)** | behavior | WTB | C4 C5 C1 C2 C3 | 4 | **24** |
|  |  |  | C2 C3 C1 C4 C5 | 3 |  |
|  |  |  | C2 C3 C4 C5 C1 | 3 |  |
|  |  |  | C1 C4 C2 C3 | 1 |  |
|  |  |  | C1 C2 C3 C4 C5 | 4 |  |
|  |  |  | C1 C4 C5 C2 C3 | 3 |  |
|  |  |  | C4 C5 C2 C3 C1 | 5 |  |
|  |  |  | C5 C3 C1 C2 C4 | 1 |  |
|  | imaging | SS96 x GCaMP7f | A2 C0 C1 C2 C3 C4 C5 A2 | 2 | **12** |
|  |  |  | A2 C0 C1 C4 C5 C2 C3 A2 | 2 |  |
|  |  |  | A2 C0 C2 C3 C1 C4 C5 A2 | 2 |  |
|  |  |  | A2 C0 C2 C3 C4 C5 C1 A2 | 2 |  |
|  |  |  | A2 C0 C4 C5 C1 C2 C3 A2 | 2 |  |
|  |  |  | A2 C0 C4 C5 C2 C3 C1 A2 | 2 |  |
| **Gradient with grass (2D)** | imaging | SS96 x GCaMP7f | A2s pIC2 sgC2 dgC2 | 2 | **12** |
|  |  |  | A2s sgC2 dgC2 pIC2 | 2 |  |
|  |  |  | A2s dgC2 pIC2 sgC2 | 2 |  |
|  |  |  | A2s dgC2 sgC2 pIC2 | 2 |  |
|  |  |  | A2s pIC2 dgC2 sgC2 | 2 |  |
|  |  |  | A2s sgC2 pIC2 dgC2 | 2 |  |
| Sun disk with grass (2D) | imaging | SS96 x GCaMP7f | A2s pIB2s sgB2s dgB2s | 2 | 12 |
|  |  |  | A2s sgB2s dgB2s pIB2s | 2 |  |
|  |  |  | A2s dgB2s pIB2s sgB2s | 2 |  |
|  |  |  | A2s dgB2s sgB2s pIB2s | 2 |  |
|  |  |  | A2s pIB2s dgB2s sgB2s | 2 |  |
|  |  |  | A2s sgB2s pIB2s dgB2s | 2 |  |

### Data analysis

All data analysis was done in Python (version 3.8). The analysis code was built on two packages that were developed as part of this study: fly2p for analysis of imaging data (https://github.com/hjmh/fly2p), and unityvr for analysis of behavior and imaging (https://github.com/hjmh/unityvr). Fly2p uses Napari (contributors, 2019) for visualizing multidimensional arrays and drawing of masks and regions of interest.

#### Preprocessing of virtual trajectories

The position and orientation of the fly in the VR environment was logged by Unity for each frame, that is, each new rendered image (behavior rig: 120 Hz, imaging rig: 144 Hz) in the form of a json log file. Using functionality implemented in the unityvr package, the log files were parsed to extract metadata and positional information, which was collected in a Pandas (version 1.2.0) data frame. These data frames were then used to compute derived parameters, such as the fly’s translational and angular velocity and the cumulative distance traveled by the fly. Occasionally, flies “jumped” and lost contact with the ball, allowing the ball to rotate freely. Since the jumps were rare and short in duration, we chose to detect and mask the position of the fly during the jump interval rather than discarding the entire trial. Jumps were found by relying on the observation that the translational velocity of a walking fly oscillates due to the characteristic gait of the fly on the ball, while the freely rotating ball does not. This effect was previously described in crickets (Haberkern and Hedwig, 2016). We computed the Fourier transform of the translational velocity and used a threshold on the power in a small frequency band (1-2 Hz) to generate a mask for “jump segments”. The trajectory data, along with metadata about the experiment and object locations in the virtual world, was saved for later analysis.

#### Analysis of virtual trajectories

##### Shape transformation

For some analyses, we reparametrized the virtual trajectories by their respective path lengths, instead of by time, so that consecutive pairs of x and y coordinates along the path were separated by a constant path length, as described previously (Haberkern et al., 2019). We chose the length increment such that the reparametrized paths had the same number of points as the original time-parameterized paths. These transformed trajectories were used for analyses aimed at quantifying trajectory shape rather than velocities, such as when we quantified tortuosity and fixation direction.

##### Computing path tortuosity

The path tortuosity of a trajectory is defined as C/L where C is the total path length of the trajectory and L is the magnitude of net displacement. The path tortuosity was log-scaled for all subsequent analysis since the values spanned several orders of magnitude.

##### Computing fixation direction

For a given trial, we estimated the direction and strength of fixation behavior as described previously (Haberkern et al., 2019). Briefly, we first computed the distribution of heading angles using 20° wide bins, and then fit the distribution with a von Mises function. The von Mises distribution has two parameters: the location parameter *μ* ∈ *[-π, π]*, which describes the location of the single peak, and the shape parameter κ, which governs the steepness of this peak. For each fit, we verified that the von Mises function approximated the heading distribution well (p>0.1, Kolmogorov Smirnov test). In some cases, flies fixated particularly strongly in a single direction leading to a sharply peaked distribution that is not well fit by a von Mises function (p<=0.1, Kolmogorov Smirnov test). If this was the case and the population vector average of the heading angles was greater than 0.5, we accepted the μ and κ despite the bad fit. Otherwise, we classified the trial as showing no fixation behavior and excluded it for analyses involving μ.

##### Preprocessing of imaging data

ScanImage saves imaging data as TIFF files, which were preprocessed in Python 3 using functions from the fly2p package. To analyze compass (EPG) neurons in the EB, we converted the 3D volume collected for each time point into a 2D image using a maximum intensity projection, resulting in a 3D image stack (x/y positions and time frames) for each trial. This image stack was then spatially filtered with a two-pixel-wide Gaussian filter and x/y motion corrected using phase correlation to a reference image. The reference image was created by averaging a small subset of frames, which were selected semi-automatically to make sure that the image was not dominated by transient activity peaks. We then performed background subtraction by selecting a region outside of the EB that did not show fluorescence changes related to calcium activity and subtracting the mean fluorescence in that region from each pixel. To compute fluorescence changes as *ΔF/F* = (*F* – *F*_0_)/*F*_0_, the fluorescence baseline *F*_0_trials where kappa was estimated by averaging over the 10% of lowest-intensity frames in each trial. We then calculated *ΔF/F* for each pixel over the whole time series, obtaining a 3D *ΔF/F* image stack. Finally, we defined regions of interest (ROIs) that covered the EB like pizza slices, approximating the arborization pattern of EPG neurons. To do this, we first defined an ellipse that covered the EB by manually marking the EB center and the long and short axis of the ellipse on the reference image. From these landmarks, an ellipse was constructed and segmented into 32 equiangular ROIs. For each ROI we computed the mean *ΔF/F* as a function of time. This time series as well as the ROI masks and reference images were saved for later analysis.

##### Calculation of bump offset

To characterize the relationship between the fly’s angular orientation within a given scene *α_worid_*, and the angular position of the bump along the circumference of the EB *α_EB_* (**Figure 2A-C**), we compute the offset ∈[–*π,π*]. The offset is simply the circular difference between *α_world_* and *α_EB_* (**Figure 1C**). We define *α_EB_* ∈ [–*π, π*] with 0 at the top of the EB as shown in **Figure 1B**. Previously, *α_EB_* was obtained for each time frame as the population vector average across ROIs (Kim et al., 2019; Seelig and Jayaraman, 2015). However, this method fails to accurately describe the position of the calcium bump when multiple activity bumps are present simultaneously, which occasionally occurs in complex visual worlds. A more robust offset measure can be calculated in the following manner, as recently proposed (Dan et al., 2021): For each frame, we detected peaks in the ΔF/F distribution across ROIs (**Figure 1D, E**). To avoid edge effects when performing peak detection on circular data, we padded array by extending it by half a phase on each side. For each peak within the original data range, we calculated the offset (**Figure 1F, G**). Thus, we could get multiple peaks or no peaks per frame. To obtain a time series of offsets, we next classified the per-frame offsets into groups that we considered to belong to a common time series. For this purpose, we obtained candidate trial offsets as the peaks in the overall distribution of peak-derived offsets. The peaks were detected on the kernel density estimation (KDE) of all per-frame offset values in a trial. We allowed up to 3 offset groups per trial. The per-frame offsets were then assigned to one of the candidate offsets based on their proximity. Finally, for each frame a main offset was chosen as the one corresponding to the largest peak (**Figure 1F, G**). For a trial, we used either the main offset time series (a compound of up to three offsets) or the most used single offset time series (primary offset), to compute the trial offset location as the circular mean and offset stability as the trial offset variance (1-Population vector average of all used offset angles).

##### Data selection criteria

We used the following criteria to select data for quantitative analysis and visualization from the original dataset on a per-trial basis: In any analysis involving the primary offset as opposed to the main offset, we only selected trials where this primary offset was used >50% of the time. This typically affected fewer than two trials per experimental group. For analysis of the offset location (circular mean), we only considered trials in which the offset variance was <0.6, to ensure that the positional readout was meaningful. For behavioral analysis of the fixation direction μ, we only considered trials where kappa was κ>0.3, to select for flies that showed at least some fixation.

### Statistics

For **Figure 3B**: ANOVA of the circular variance of primary offset (min. 50% usage) across gradient trials, comparing the primed and unprimed group.

|  | sum_sq | df | F | PR(>F) |
| --- | --- | --- | --- | --- |
| C(stimulus) | 1.0995713728713700 | 3.0 | 7.772609802215830 | **8.71112756687187E-05** |
| C(group) | 0.14878717006825000 | 1.0 | 3.155223876454220 | **0.07821854517040300** |
| C(stimulus):C(group) | 0.09838483196746690 | 3.0 | 0.6954590927033410 | 0.5565749761694800 |
| Residual | 5.6586984338665900 | 120.0 |  |  |

For **Figure 3C**: Rayleigh test on filtered primary offset values for deviation from uniformity.

| Stimulus | Sun | Low gradient | Very low gradient | Low bright gradient |
| --- | --- | --- | --- | --- |
| Primed | p = 0.0009, *** | p = 0.0110, * | p = 0.1357, ns | p = 0.0633, ns |
| Unprimed | n/a | p = 0.0009, *** | Not enough data | p = 0.1357, ns |

For **Figure 3C:** No. of trials excluded per group based on whether flies had a stable offset primary offset variance <0.6)

| Group | Stimulus | Excluded |
| --- | --- | --- |
| Unprimed | C1t01 | 2 |
| Unprimed | C2t01 | 5 |
| Unprimed | C3t01 | 10 |
| Unprimed | C4t01 | 4 |
| Primed | C1t01 | 3 |
| Primed | C2t01 | 5 |
| Primed | C3t01 | 6 |
| Primed | C4t01 | 1 |

**For Figure 3D:** No. of trials excluded per group based on whether flies showed fixation unimodal fixation with kappa>0.2) and had a stable offset (primary offset variance <0.6).

| Group | Stimulus | Excluded |
| --- | --- | --- |
| Unprimed | C1t01 | 3 |
| Unprimed | C2t01 | 7 |
| Unprimed | C3t01 | 11 |
| Unprimed | C4t01 | 6 |
| Primed | A2t01 | 3 |
| Primed | C1t01 | 5 |
| Primed | C2t01 | 6 |
| Primed | C3t01 | 8 |
| Primed | C4t01 | 1 |

**For Figure 3E:** ANOVA on log(tortuosity) values across gradient trials comparing primed and unprimed groups in the imaging dataset

|  | sum_sq | df | F | PR(>F) |
| --- | --- | --- | --- | --- |
| Group | 3.537530592070450 | 1.0 | 3.5946668310008900 | 0.060370106072757900 |
| Dtimulus | 6.6261298392209600 | 3.0 | 2.2443838087076600 | 0.0866323027366014 |
| Group:Stimulus | 0.4036066958217400 | 3.0 | 0.13670850936642900 | 0.9379346888497690 |
| Residual | 118.09263306058800 | 120.0 |  |  |

**For Figure 3H**: Rayleigh test on filtered mu values for deviation from uniformity.

| Stimulus | Group | N | Excluded | rayleigh_p |  |
| --- | --- | --- | --- | --- | --- |
| Sun | primed | 16 | 3.0 | 0.11542426476831700 | ns |
| Gradient | unprimed | 16 | 1.0 | 0.03468572395017710 | *p<0.05 |
| Gradient | primed | 16 | 3.0 | 0.1622115186543340 | ns |
| Low gradient | unprimed | 16 | 4.0 | 0.011676623894337600 | *p<0.05 |
| Low gradient | primed | 16 | 1.0 | 0.4204615996453210 | ns |
| Very low gradient | unprimed | 16 | 5.0 | 0.19738995914062400 | ns |
| Very low gradient | primed | 16 | 4.0 | 0.5365459728272820 | ns |
| Bright low gradient | unprimed | 16 | 3.0 | 0.03153848622535020 | *p<0.05 |
| Bright low gradient | primed | 16 |  | 0.4266437940955400 | ns |

For **Figure 3 Supplement 3B**: ANOVA of the log(tortuosity) values across gradient trials (stimuli), comparing primed and unprimed group in the behavior setup

|  | sum_sq | df | F | PR(>F) |
| --- | --- | --- | --- | --- |
| Group | 0.7159924308133220 | 1.0 | 1.9270608685180300 | 0.1671115933244710 |
| Stimulus | 13.03612418405390 | 3.0 | 11.69537726522960 | 6.14095919734344E-07 |
| Group:Stimulus | 0.1637136416015870 | 3.0 | 0.14687592531047300 | 0.9315563960156730 |
| Residual | 56.47504511225900 | 152.0 |  |  |

For **Figure 3 Supplement 3A**: Rayleigh test for uniform circular distribution.

| stimulus | group | N | excluded | rayleigh_p |  |
| --- | --- | --- | --- | --- | --- |
| Sun | Primed (imaging) | 16 | 3.0 | 0.11542426476831700 | ns |
| Sun | Primed (behavior) | 20 | 3.0 | 0.5713014916995520 | ns |
| Gradient | Unprimed (imaging) | 16 | 1.0 | 0.03468572395017710 | p<0.05 |
| Gradient | Primed (imaging) | 16 | 3.0 | 0.1622115186543340 | ns |
| Gradient | Unprimed (behavior) | 20 | 1.0 | 4.26411325151675E-07 | p<0.001 |
| Gradient | Primed (behavior) | 20 |  | 1.56330792892864E-07 | p<0.001 |
| Low gradient | Unprimed (imaging) | 16 | 4.0 | 0.011676623894337600 | p<0.05 |
| Low gradient | Primed (imaging) | 16 | 1.0 | 0.4204615996453210 | ns |
| Low gradient | Unprimed (behavior) | 20 | 2.0 | 7.83508009789455E-07 | p<0.001 |
| Low gradient | Primed (behavior) | 20 | 3.0 | 0.0029396855129062900 | p<0.01 |
| Very low gradient | Unprimed (imaging) | 16 | 5.0 | 0.19738995914062400 | ns |
| Very low gradient | Primed (imaging) | 16 | 4.0 | 0.5365459728272820 | ns |
| Very low gradient | Unprimed (behavior) | 20 | 6.0 | 5.17726537781011E-05 | p<0.001 |
| Very low gradient | Primed (behavior) | 20 | 7.0 | 0.5817427141412790 | ns |
| Low bright gradient | Unprimed (imaging) | 16 | 3.0 | 0.03153848622535020 | p<0.05 |
| Low bright gradient | Primed (imaging) | 16 |  | 0.4266437940955400 | ns |
| Low bright gradient | Unprimed (behavior) | 20 | 5.0 | 0.0006503499262968700 | p<0.001 |
| Low bright gradient | Primed (behavior) | 20 | 4.0 | 0.7333454469496580 | ns |

For **Figure 4D**: ANOVA of circular variance of offset 1 (min. 50% usage) across single and combined stimulus trials.

|  | sum_sq | df | F | PR(>F) |
| --- | --- | --- | --- | --- |
| **C(Stimulus)** | 0.31425895015750300 | 4.0 | 2.7832988946266900 | **0.030146419141029800** |
| **C(Group)** | 0.02465670100175980 | 1.0 | 0.8735085331245910 | 0.3520329937575080 |
| **C(Stimulus):C(Group)** | 0.10021577732117900 | 4.0 | 0.8875816014226360 | 0.4739701727293970 |
| **Residual** | 3.1049921178122200 | 110.0 |  |  |

For **Figure 4E**: ANOVA of log(tortuosity) across single and combined stimulus trials comparing naturalistic and conflicting scenes on the imaging dataset

|  | sum_sq | df | F | PR(>F) |
| --- | --- | --- | --- | --- |
| **Group** | 5.647993029797080 | 1.0 | 9.296888517014170 | 0.0029376955089513400 |
| **Stimulus** | 3.7569869747920000 | 4.0 | 1.5460486973091400 | 0.19471812035991700 |
| **Group:Stimulus** | 0.915448994154827 | 4.0 | 0.37671909281623000 | 0.8247800093937010 |
| **Residual** | 60.751433336655900 | 100.0 |  |  |

For **Figure 4 Supplement 1D**: ANOVA of circular variance of offset 1 (min. 50% usage) between first and second sun stimulus trial across the two groups.

|  | sum_sq | df | F | PR(>F) |
| --- | --- | --- | --- | --- |
| **C(pre/post)** | 0.04726691608282040 | 1.0 | 1.257146387101270 | 0.2692276596215800 |
| **C(group)** | 2.11169105405376E-05 | 1.0 | 0.000561641206848828 | 0.9812167408078310 |
| **C(pre/post):C(group)** | 0.0001379581313887820 | 1.0 | 0.003669238038351420 | 0.9520157586472780 |
| **Residual** | 1.4287459516061000 | 38.0 |  |  |

For **Figure 4 Supplement 2C**: P-values for Rayleigh test on fixation direction normalized to the fixation direction in the low gradient for behavior and imaging data. Trials were excluded if the flies were not fixating (unimodal fixation and kappa<0.2)

| Stimulus | Group | N | Excluded | Rayleigh p value |
| --- | --- | --- | --- | --- |
| Sun (pre) | Naturalistic (imaging) | 12 | 1.0 | 0.13270147233266400 |
| Sun (pre) | Conflict (imaging) | 12 |  | 0.5807612230832790 |
| Sun (pre) | Naturalistic (behavior) | 20 | 1.0 | 0.04170527257336140 |
| Sun (pre) | Conflict (behavior) | 20 | 2.0 | 0.9572962504864780 |
| Sun (post) | Naturalistic (imaging) | 12 | 1.0 | 0.00034101821682623300 |
| Sun (post) | Conflict (imaging) | 12 | 1.0 | 0.43512341840575900 |
| Sun (post) | Naturalistic (behavior) | 20 | 4.0 | 0.13749277832941200 |
| Sun (post) | Conflict (behavior) | 20 | 2.0 | 0.31817975958811600 |
| Low gradient | Naturalistic (imaging) | 12 | 1.0 | n/a |
| Low gradient | Conflict (imaging) | 12 |  | n/a |
| Low gradient | Naturalistic (behavior) | 20 | 4.0 | n/a |
| Low gradient | Conflict (behavior) | 20 | 5.0 | n/a |
| Natural combi1 | Naturalistic (imaging) | 12 | 1.0 | 0.01513639418049750 |
| Conflict combi1 | Conflict (imaging) | 12 |  | 0.1425965280962570 |
| Natural combi1 | Naturalistic (behavior) | 20 | 1.0 | 0.0027574525648617200 |
| Conflict combi1 | Conflict (behavior) | 20 | 4.0 | 0.48051216718781000 |
| Natural combi2 | Naturalistic (imaging) | 12 |  | 0.020393252231252900 |
| Conflict combi2 | Conflict (imaging) | 12 | 1.0 | 0.03365709811919680 |
| Natural combi2 | Naturalistic (behavior) | 20 | 1.0 | 0.011447747236754500 |
| Conflict combi2 | Conflict (behavior) | 20 | 2.0 | 0.035739597202308100 |

For **Figure 4 Supplement 2D**: ANOVA of log(tortuosity) across single and combined stimulus trials comparing naturalistic and conflicting scenes on the behavior dataset

|  | sum_sq | df | F | PR(>F) |
| --- | --- | --- | --- | --- |
| **Group** | 0.6617876543291180 | 1.0 | 1.3255570312360100 | 0.2510439356542950 |
| **Stimulus** | 3.3265486211982000 | 4.0 | 1.6657646473655800 | 0.15962508469941200 |
| **Group:Stimulus** | 0.5970524110138840 | 4.0 | 0.2989731737433860 | 0.8783620467209530 |
| **Residual** | 94.85797393816130 | 190.0 |  |  |

For **Figure 5 C**: ANOVA for the primary offset variance using the grass density (“density”) and the sky texture (“group”) as well as their interaction as explanatory variables.

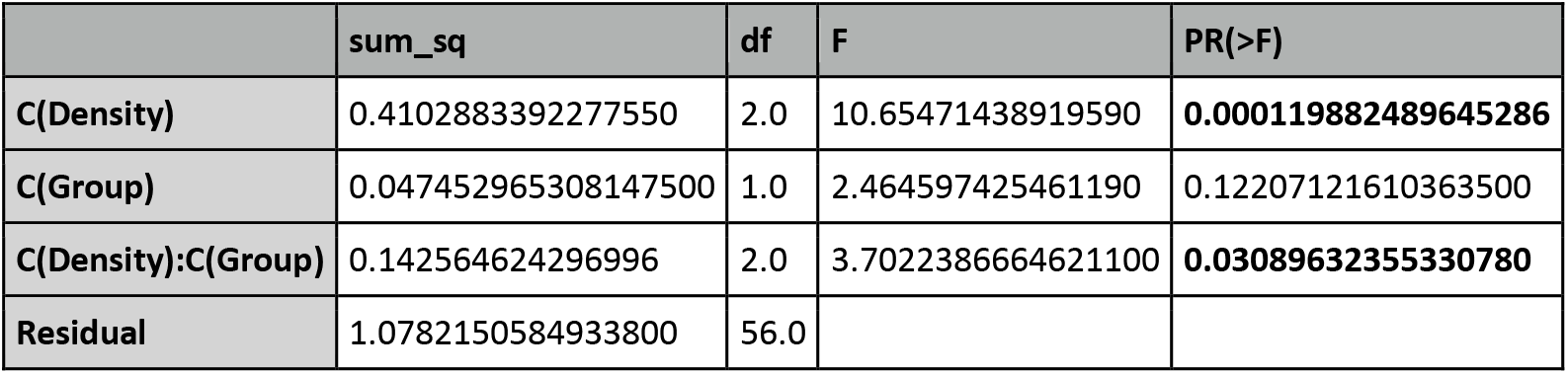

---

**Figure 1 Supplement 1: Display dimensions.**

**A:** Top view of the pentagonal display used in the imaging rig and its relation to the treadmill.

**B:** Side view of the screen and trackball in the imaging rig, illustrating occlusion of the display by the trackball and the fly holder.

**C:** Top view of the V-shaped display used in the behavior rig.

**D:** Side view of the screen and trackball in the behavior rig

**Figure 1 Supplement 2: Overview of visual panoramas used as distal cues.**

**A:** Sun disk panoramas. Ai: Panorama images with names and azimuthal position convention. Aii: Intensity profile along the vertical panorama axis at +/-π, illustrating the position of the sun disk.

**B:** Gradient panoramas. Bi: Panorama images with names and azimuthal position convention. Bii: Intensity profile of gradient panoramas along the azimuthal position.

**C:** Panoramas that combine a sun disk and gradient. Cii: Sun disk and gradient panoramas in naturalistic configuration (top two) and in conflicting configuration (bottom two). Cii: Intensity profile of sun disk and gradient panoramas along the azimuthal position.

**Figure 1 Supplement 3: Display of visual stimuli on the different screen layouts.**

**A:** Definition of the horizon line and coverage of the screen by the simulated cylinder in (1D) panorama experiments. In the imaging rig, the fly is positioned closer to the top edge of the screen and this the horizon line lies above the midline in elevation. The lower part of the screen is covered by the treadmill in this configuration and therefore not visible to the fly. To lower imaging noise, a black ground plane is inserted in these cases. The upper field of view is occluded by the imaging holder. The screen in the behavior rig covers a larger elevation above the horizon line as the fly’s field of view is not restricted by a holder. The lower field of view is limited by the ball, just like in the imaging case.

**B:** Illustration of how the panorama textures are mapped onto a virtual cylinder to simulate distal panoramic environments.

**C:** Reference frames in relation to the screen geometry.

**D:** Illustration of the mapping of stimuli in two-dimensional worlds onto the faces of the panoramic screen. Left: Schematic of what is visible at different elevations. Right: Screenshot of a frame in a two-dimensional world with local objects (inset shows the ground plane texture).

**Figure 1 Supplement 4: Brightness correction for projector-based displays.**

**A:** Gamma curve for projector display in the behavior rig. Black dots show the measured pixel values (using a power meter) as a function of specified pixel values. The green line shows the interpolated function *(f)* representing the gamma transformation for the display. The interpolated function f is used to reshape images to correct for the measured gamma curve.

**B.** Same as A but for the imaging rig. The interpolated function *(g)* is plotted in violet.

**C.** Intensity profile for the desired (black), simulated (blue, *g* transformed), and reshaped (orange, *g^-1^* transformed) gradient image on the imaging rig. The blue curve shows the *g* transformed intensity gradient and represents the profile as it would appear if the sinusoid was projected as is on the display. The orange curve shows the *g^-1^* transformed reshaped gradient. When projected on the display, the reshaped gradient will appear as intended.

**D.** Panoramic intensity gradients (center) are reshaped using *f^-1^* so that they appear as intended on the behavior rig (left) and using *g^-1^* to appear as intended on the imaging rig (right).

**Figure 1 Supplement 5: Fitting a von Mises function to the head direction distribution.**

**A:** A sample heading distribution for a fly and the fitted von-Mises function with an illustration of the interpretation of the parameters μ and κ of a von-Mises distribution: μ captures the direction of fixation for the fly, whereas κ captures the magnitude or strength of fixation.

**B:** Polar fixation plot visualizing the two fitted parameters. The fixation direction, μ, is plotted on the angular axis, and κ is plotted on the radial axis.

**Figure 2 Supplement 1: Primary vs overall offset**

Frequency distributions of the overall and the primary offset values for three flies (SS00096 > GCaMP7, same as in Figure 2G). Solid lines show distributions based on the overall offset, dashed lines the distributions based on the primary offset. For all but one trial the distributions are identical, because only one stable offset was assumed (see **Methods** for details).

**A:** Sun disk trials.

**B:** Gradient trials.

**Figure 3 Supplement 1: Offset usage**

**A:** Number of bumps detected per trial. Left: unprimed group. Right: primed group. **B:** Mean occurrence of the primary, secondary, and tertiary offsets. Left: unprimed group. Right: primed group.

**C:** Usage of the primary offset as percent of the total trial time. Left: unprimed group. Right: primed group.

**Figure 3 Supplement 2: Extended information on offset location and variance**

**A, C:** Normalized primary offset location visualized as in Figure 2K. **A:** unprimed group. **C:** primed group.

**B, D:** Circular variance of the primary offset visualized as in Figure 2J. per fly. The black line marks the median across flies. **B:** unprimed group. **D:** primed group.

**E:** The primary offset variance and the cumulative path length are not correlated. Lines are fitted using a linear mixed effects model (p-values of fit are shown in figure).

**F:** The primary offset variance and the linear displacement are negatively correlated. Fit same as E.

**Figure 3 Supplement 3: Comparison of behavior in the imaging and behavior only groups.**

**A:** Distribution of fixation direction across flies and trials. Comparison of four groups in purely behavioral experiments (primed: light purple; unprimed: deep purple) and experiments involving calcium imaging with behavior (primed: light green; unprimed: dark green). We found that unprimed flies had fixation directions clustered around 0°. Stars represent significant deviation from a circular uniform distribution (*p<0.05, **p<0.01, Rayleigh test for deviation from uniform distribution). See **Methods** for details.

**B:** Same as A but for log scaled path tortuosity. Black bars indicate the medians of the distributions.

**Figure 4 Supplement 1: Offset usage and offset variance per fly.**

**A:** Usage of offset 1 (primary offset) as percent of the total trial time. Left: Naturalistic stimulus group. Right: Conflicting stimulus group.

**B:** Number of bumps detected per trial. Left: Naturalistic stimulus group. Right: Conflicting stimulus group.

**C:** Circular variance of offset 1 per fly. The black line marks the median across flies. Left: Naturalistic stimulus group. Right: Conflicting stimulus group.

**D:** Direct comparison of the offset variance between the initial and final sun disk trial. No difference was detected between groups, trials and their interaction (ANOVA, see Statistics for details).

**Figure 4 Supplement 2: Behavior in conditions with sun disk and gradient compound stimuli.**

**A:** Trajectories of flies in naturalistic stimulus conditions

**B:** Trajectories of flies in conflicting stimulus conditions

**C:** Plot of fixation direction normalized to the fixation direction in the low gradient (see Fig 5F and 5G) for both imaging and behavior experiments in naturalistic and conflicting conditions. Trials were excluded based on whether flies were fixating.

**D.** Path tortuosity values for naturalistic and conflicting stimuli in imaging and behavior setups.

**Figure 5 Supplement 1: Test of compass stability with different sun disk stimuli.**

Five different sun stimuli with four different sizes were tests (see **Methods**, n=16, SS96 > GCaMP7f). The order of stimulus presentations was varied systematically to account for memory effects.

**A:** Primary offset after normalization to the offset used in the trial with the largest sun. **B:** Primary offset variance.

**C**: The percent of the trial time during which the primary offset was used. In conditions with a more challenging stimulus (smaller sun disk or less contrast), additional offsets start to form.

**Figure 5 Supplement 2: Extended information on compass stability in cluttered environments.**

**A:** Accumulated offset usage across grass densities and experimental groups (sun sky in yellow and gradient in blue). Black bars mark the median. In the dense grass condition flies in the sun sky group have a higher frequency of no offset at all (total offset usage below 100%).

**B:** The walking trajectory of a fly in the dense grass environment with a gradient sky, visualized as described in **Figure 5 G**. Note the high stability of the offset.

**C Top:** Calcium activity in EPG neurons in the EB and head direction (green) for the same fly as shown in **B**. **Bottom**: The primary offset for the calcium activity and heading direction shown above.

## Notes

### Competing Interest Statement

The authors have declared no competing interest.

