## Supplementary figures and images for "Maintaining a stable head direction representation in naturalistic visual environments"

### Figure1Figure Supplement 1

#### Figure 1 Supplement 1: Display dimensions

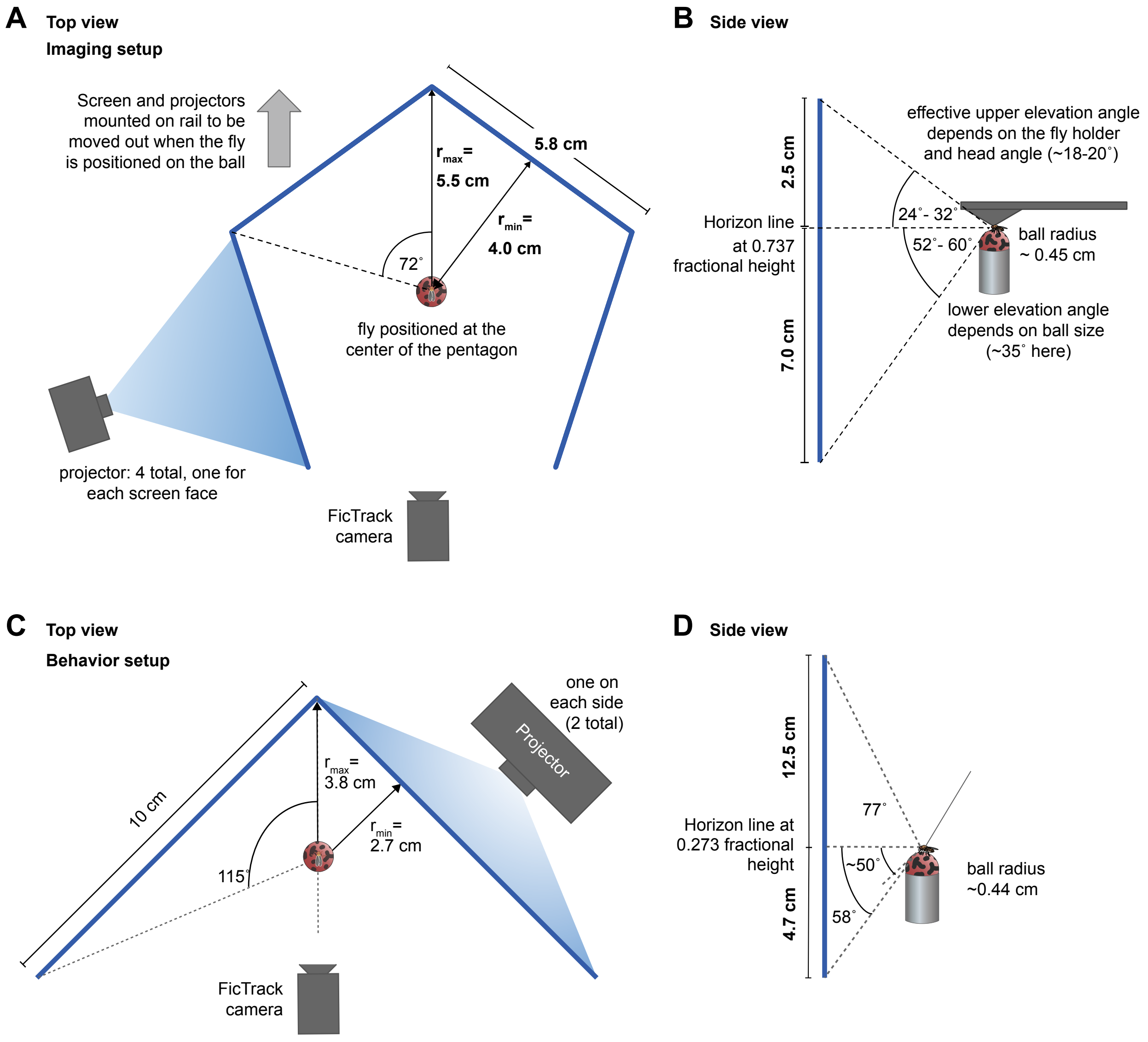

### Figure1Figure Supplement 2

#### Figure 1 Supplement 2: Overview of visual panoramas

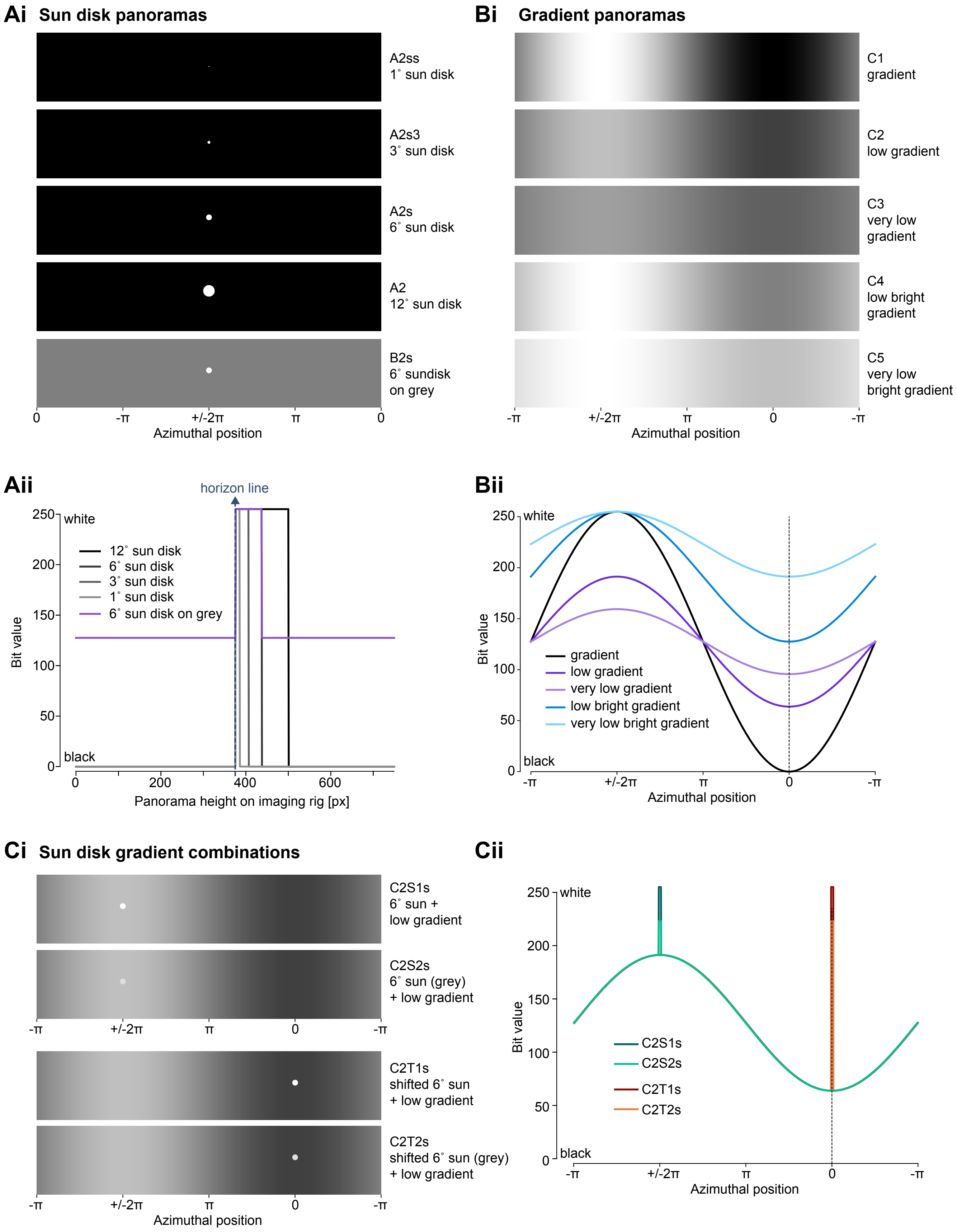

### Figure1Figure Supplement 3

#### Figure 1 Supplement 3: Display of visual stimuli on two different screens

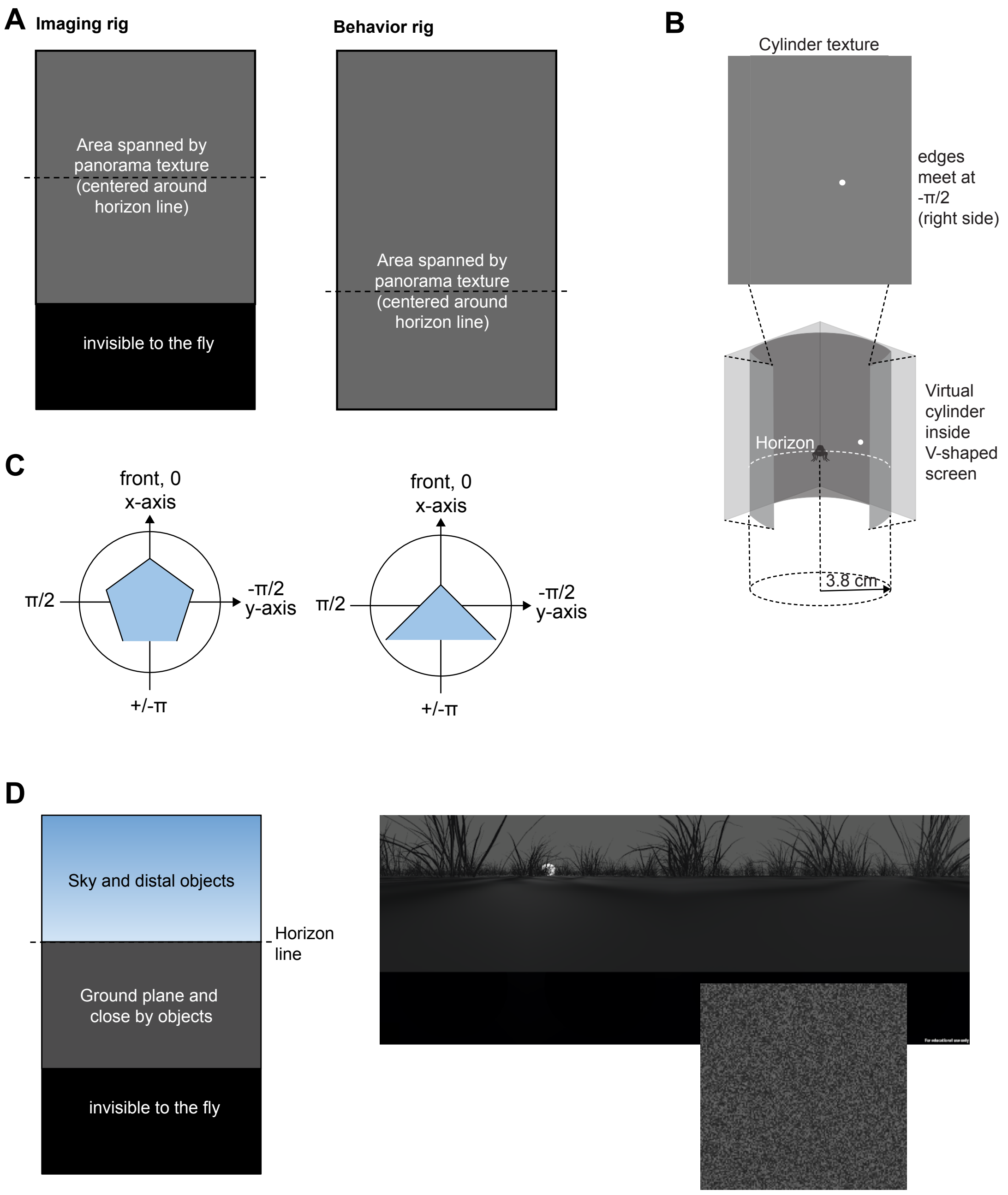

### Figure1Figure Supplement 4

#### Figure 1 Supplement 4: Brightness correction for projector-based display

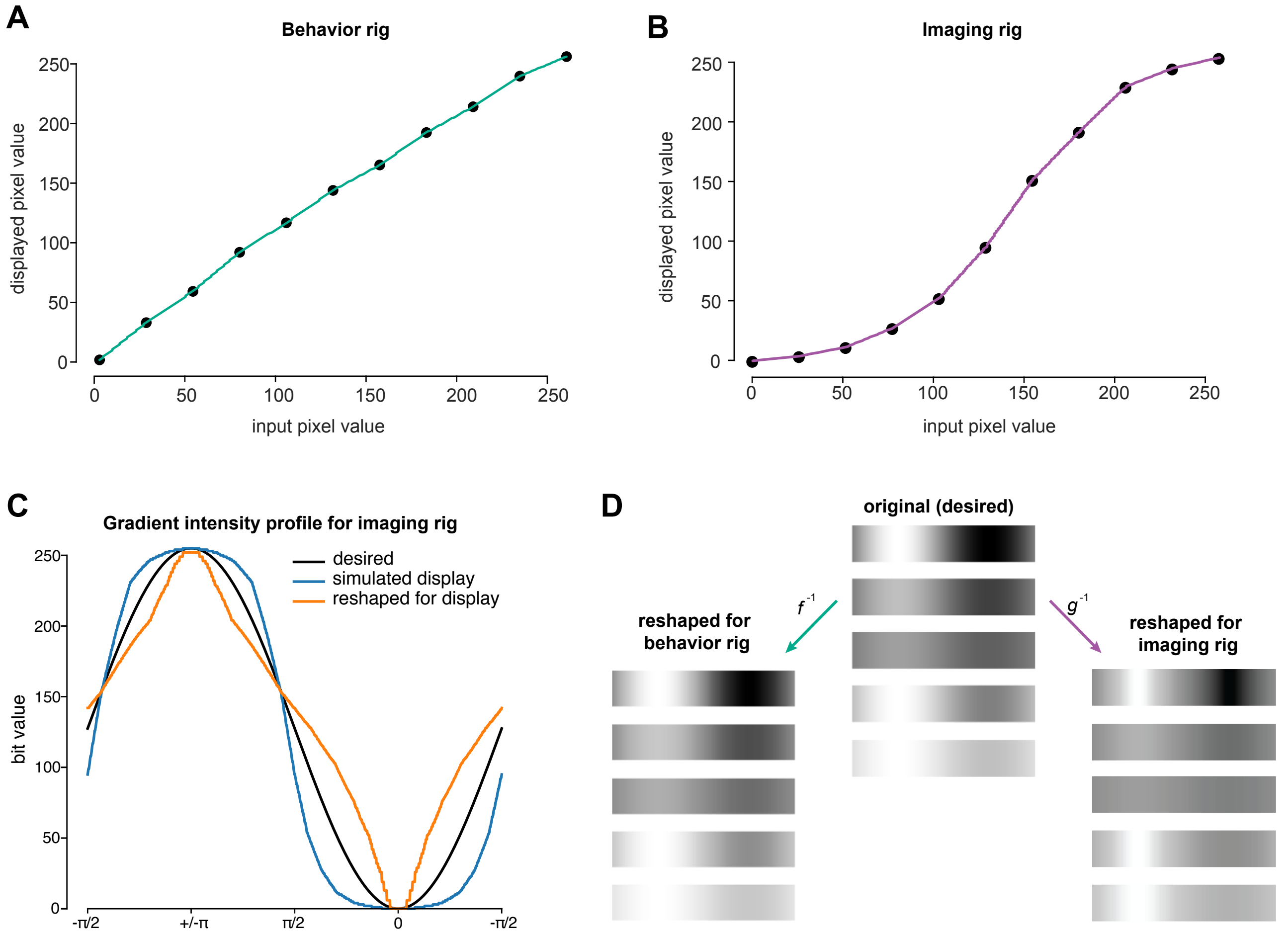

### Figure1Figure Supplement 5

#### Figure 1 Supplement 5: Fitting von Mises function to the head direction distribution

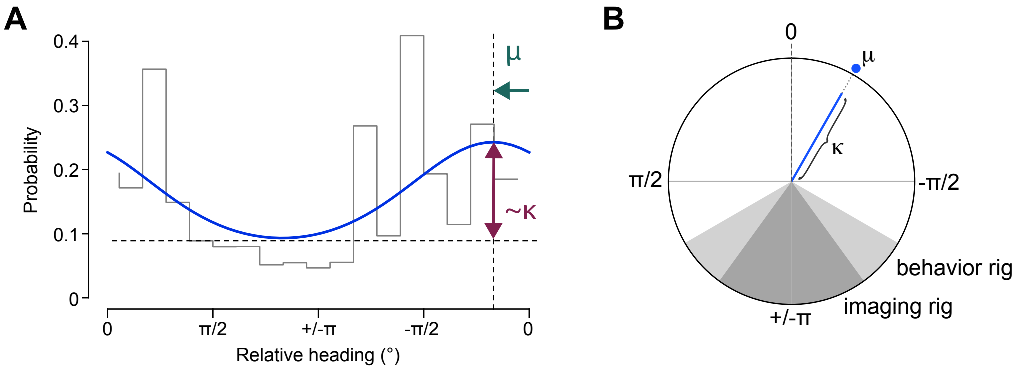

### Figure2Figure Supplement 1

##### Figure 2 Supplement 1: Primary vs overall offset

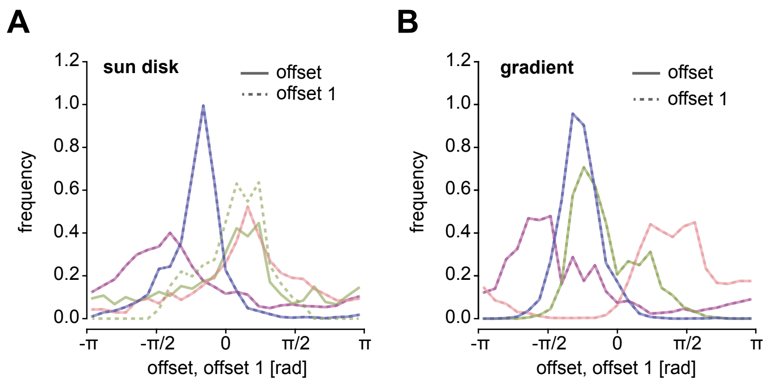

### Figure3Figure Supplement 1

#### Figure 3 Supplement 1: Offset usage

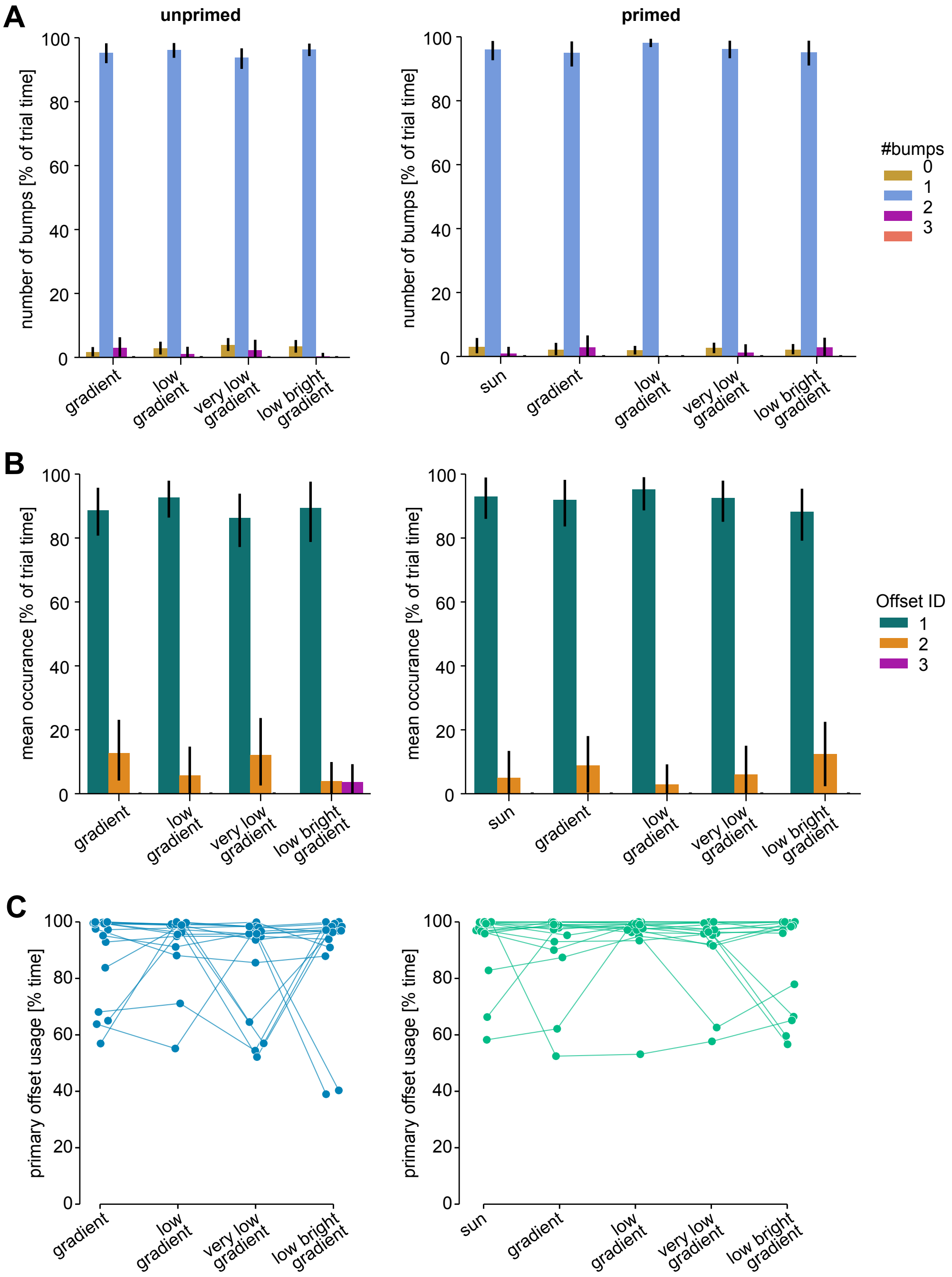

### Figure3Figure Supplement 2

#### Figure 3 Supplement 2: Extended information on offset location and variance

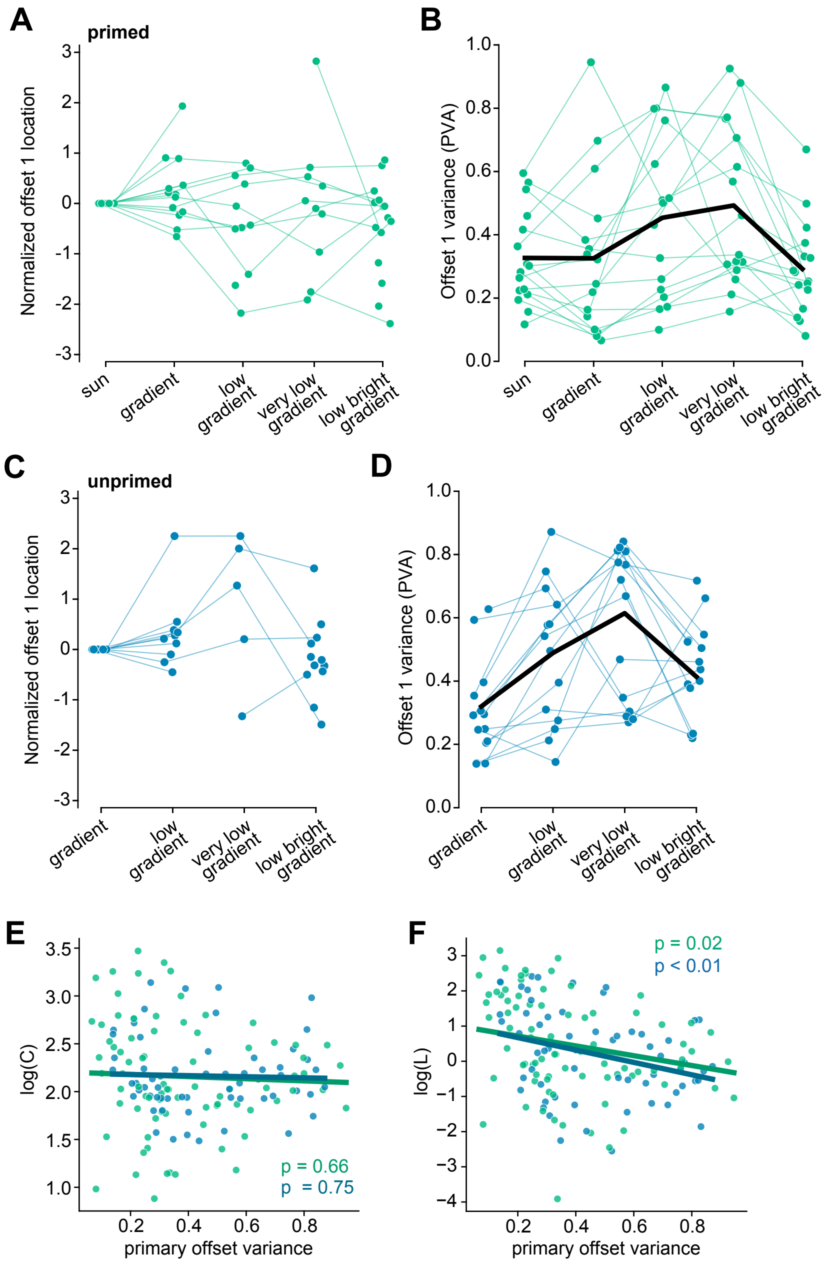

### Figure4Figure Supplement 1

##### Figure 4 Supplement 1: Offset usage and offset variance per fly

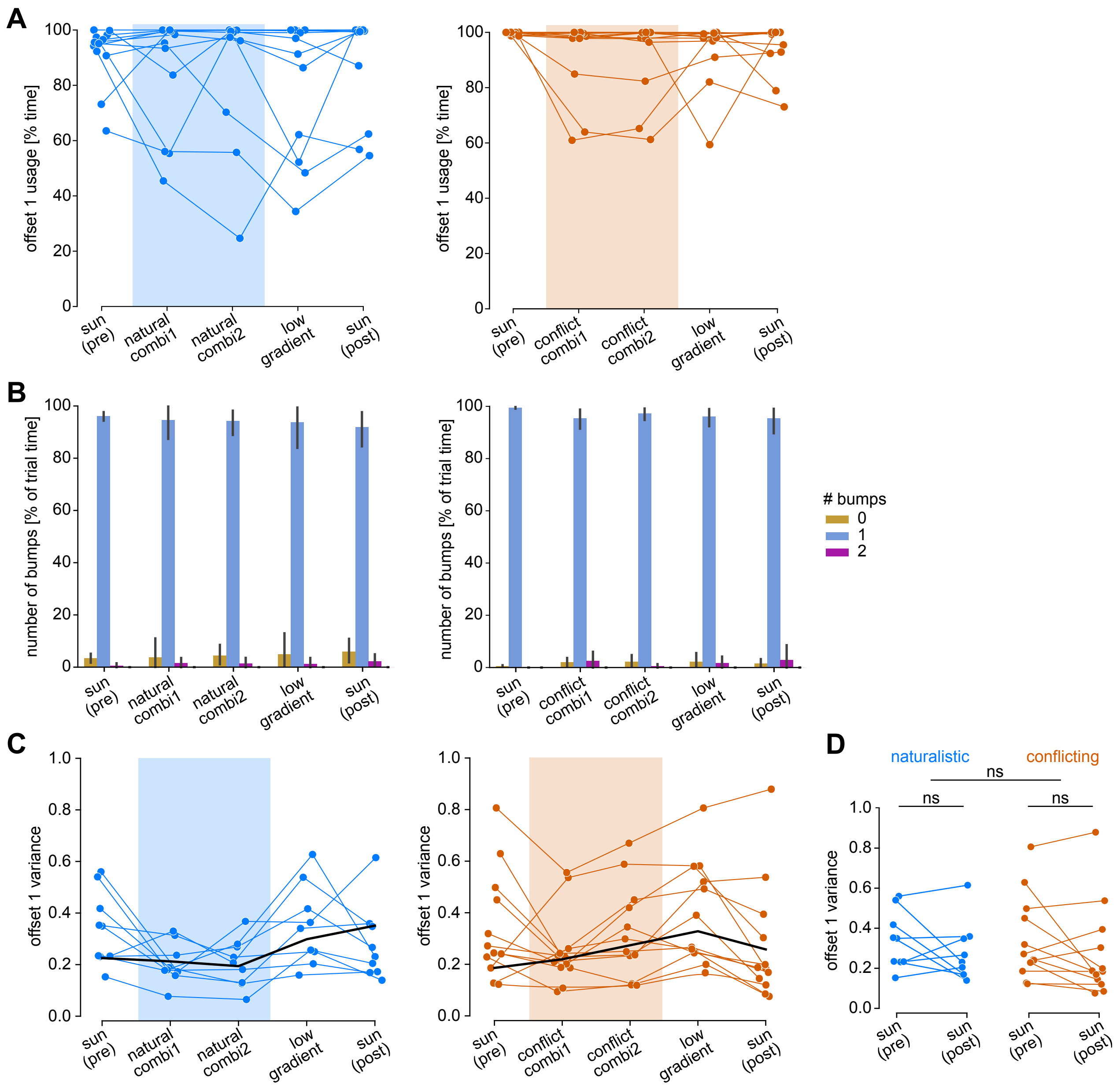

### Figure4Figure Supplement 2

**Figure 4 Supplement 2: Behavior in conditions with sun disk and gradient compound stimuli.**

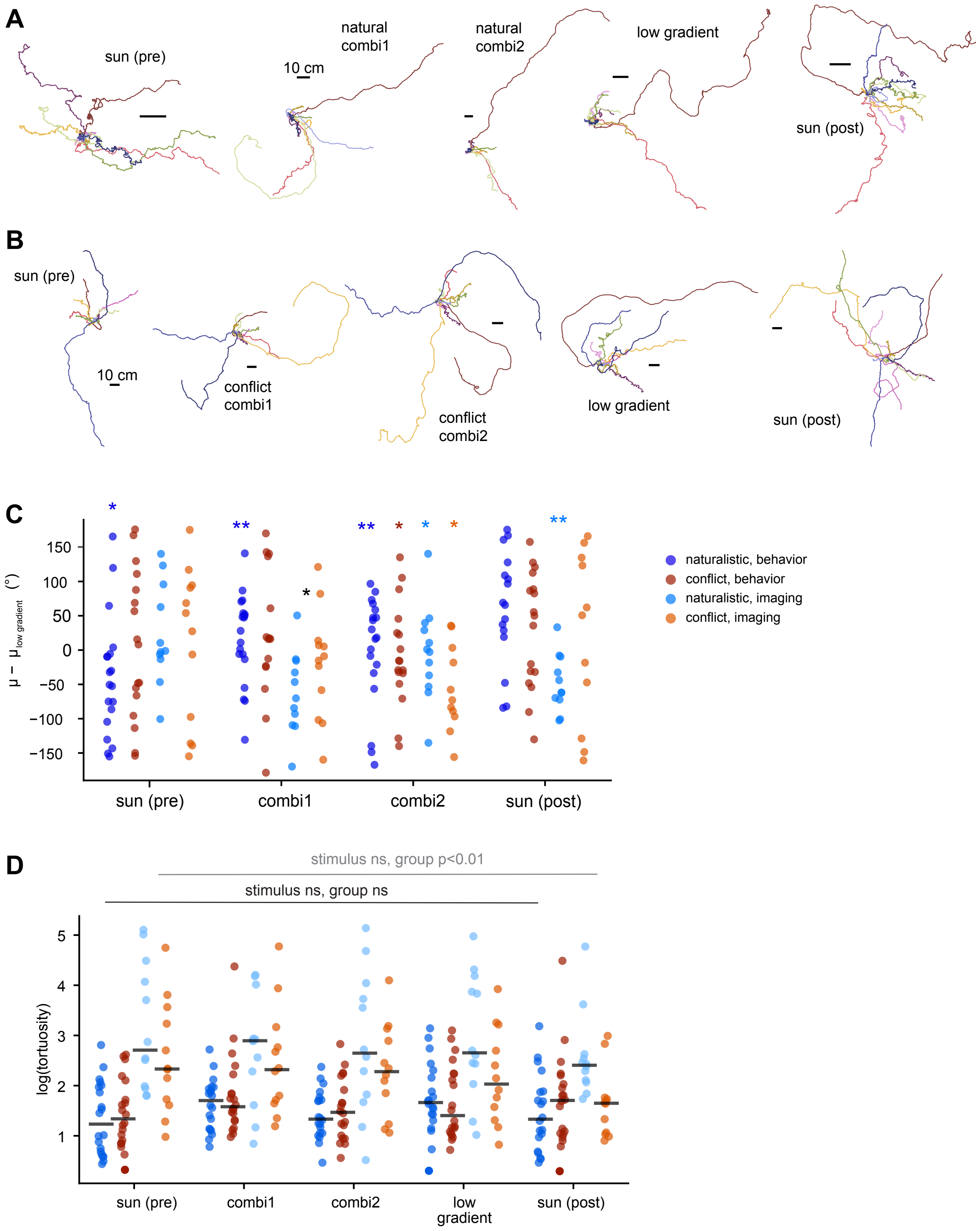

### Figure5Figure Supplement 1

##### Figure 5 Supplement 1: Test of compass stability with different sun disk stimuli

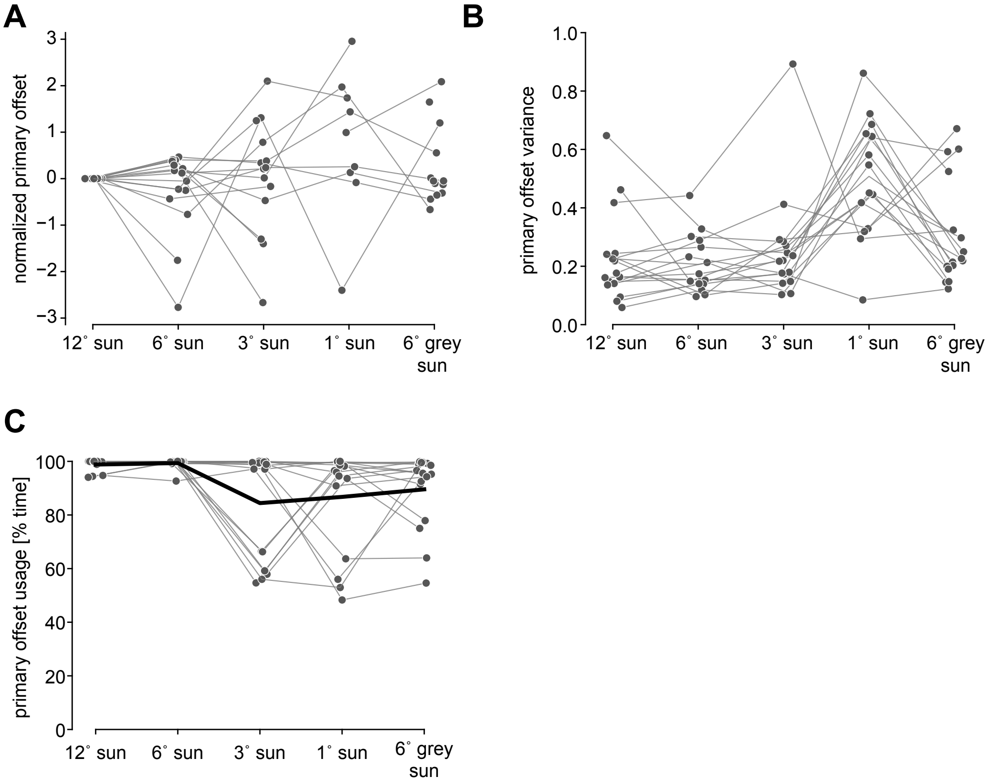

### Figure5Figure Supplement 2

##### Figure 5 Supplement 2: Extended information on compass stability in cluttered environments.

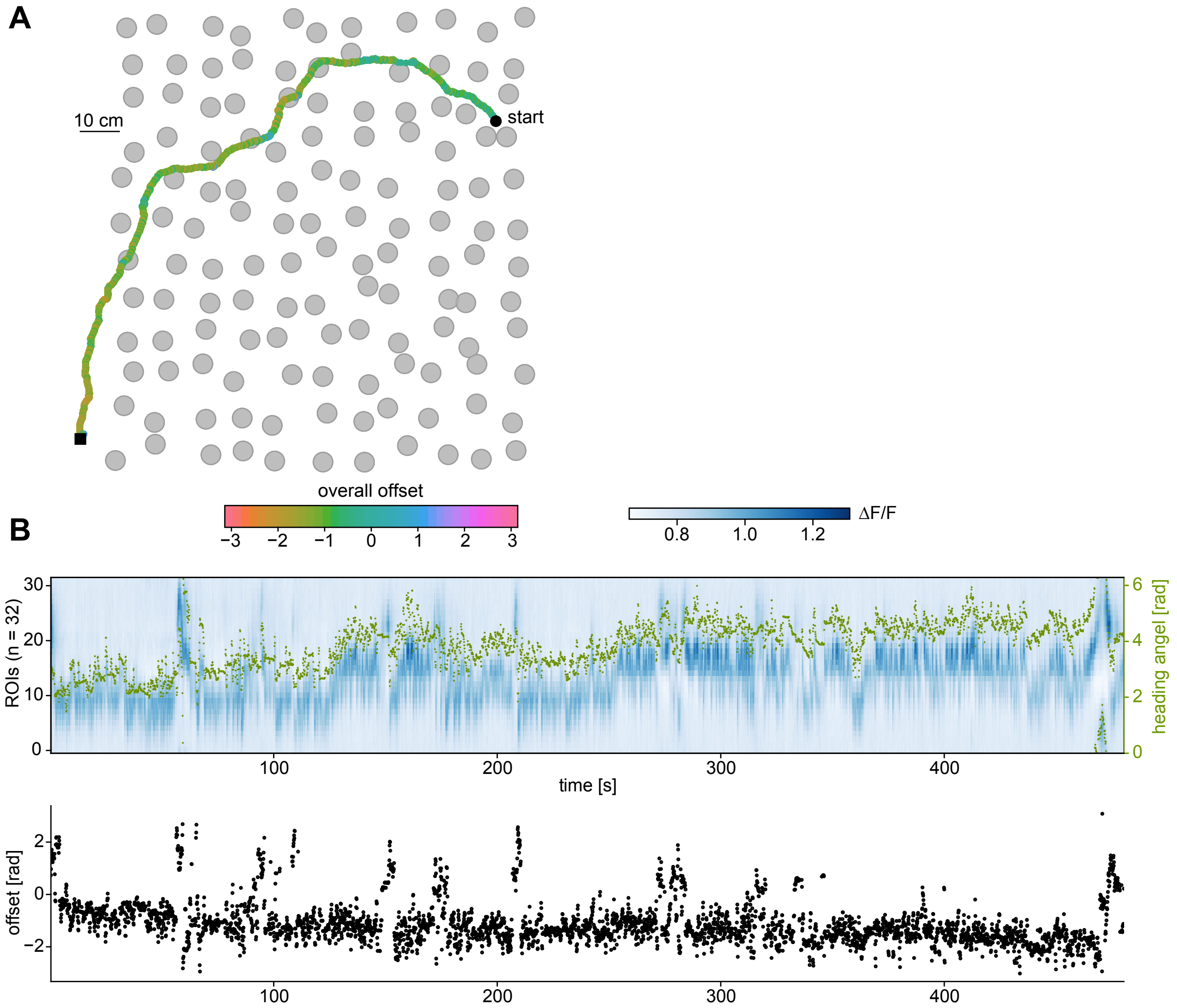

### Supplemental Data 1

#### Figure 3 Supplement 3: Comparison of behavior in the imaging and behavior only groups

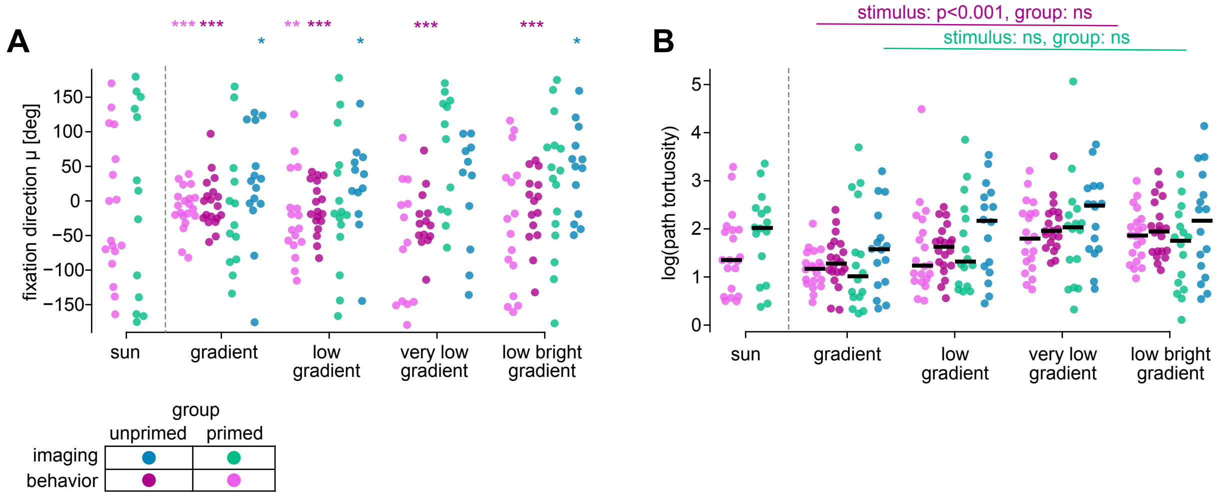
